# Scalable, accessible, and reproducible reference genome assembly and evaluation in Galaxy

**DOI:** 10.1101/2023.06.28.546576

**Authors:** Delphine Larivière, Linelle Abueg, Nadolina Brajuka, Cristóbal Gallardo-Alba, Bjorn Grüning, Byung June Ko, Alex Ostrovsky, Marc Palmada-Flores, Brandon D. Pickett, Keon Rabbani, Jennifer R. Balacco, Mark Chaisson, Haoyu Cheng, Joanna Collins, Alexandra Denisova, Olivier Fedrigo, Guido Roberto Gallo, Alice Maria Giani, Grenville MacDonald Gooder, Nivesh Jain, Cassidy Johnson, Heebal Kim, Chul Lee, Tomas Marques-Bonet, Brian O’Toole, Arang Rhie, Simona Secomandi, Marcella Sozzoni, Tatiana Tilley, Marcela Uliano-Silva, Marius van den Beek, Robert M. Waterhouse, Adam M. Phillippy, Erich D. Jarvis, Michael C. Schatz, Anton Nekrutenko, Giulio Formenti

## Abstract

Improvements in genome sequencing and assembly are enabling high-quality reference genomes for all species. However, the assembly process is still laborious, computationally and technically demanding, lacks standards for reproducibility, and is not readily scalable. Here we present the latest Vertebrate Genomes Project assembly pipeline and demonstrate that it delivers high-quality reference genomes at scale across a set of vertebrate species arising over the last ∼500 million years. The pipeline is versatile and combines PacBio HiFi long-reads and Hi-C-based haplotype phasing in a new graph-based paradigm. Standardized quality control is performed automatically to troubleshoot assembly issues and assess biological complexities. We make the pipeline freely accessible through Galaxy, accommodating researchers even without local computational resources and enhanced reproducibility by democratizing the training and assembly process. We demonstrate the flexibility and reliability of the pipeline by assembling reference genomes for 51 vertebrate species from major taxonomic groups (fish, amphibians, reptiles, birds, and mammals).

## Introduction

The vast majority of Earth’s biodiversity lacks reference genomes, leaving key questions in systematics, population biology, molecular genetics, evolution, and biogeography unanswered^1–3^. To address these questions, numerous international scientific initiatives, many collectively under the Earth BioGenome Project, aim to generate reference genomes for a wide variety of taxonomic groups, ultimately targeting all ∼1.8 million known eukaryotic species^4^. Despite these developments, cataloging the genomes from a significant fraction of biodiversity in a reasonable timeframe will require the pace of reference genome generation to increase by at least one hundredfold^1^. Achieving such speedup requires overcoming two major obstacles.

The first obstacle is developing an optimized computational approach that can be utilized by researchers at various levels of assembly expertise. Assembly involves multiple components, with software continuously evolving, driven by algorithmic improvements as well as by the advances in long-read sequencing^5, 6^. Selecting combinations of tools that generate the best results given available data types, genome sizes, heterozygosity levels, and other factors requires experience and practical knowledge of genome assembly as well as extensive testing^6^. Once determined, such best-practices need to be translated into computational workflows in order to be used by researchers beyond traditional bioinformaticians. Such workflows need to be continuously maintained and updated, adapting to the fast-evolving sequencing and assembly technology.

The second obstacle is computational infrastructure. While the hardware requirements for genome assembly have decreased substantially, work still requires highly-parallel, multi-CPU computational nodes with substantial RAM (∼0.5 TB) for an average vertebrate-sized (∼1–3 Gbp) genome, or larger for plant and animal taxa with very large (4–100 Gbp) genomes. Access to such infrastructure is limited to a few research groups, even in countries with a sustained investment in research, as it requires expertise in system design, configuration, maintenance, and large capital expenditures. Commercial cloud computing resources do not fully address these problems because these resources still need to be configured and funded. In addition, computational clouds are predominantly based in a small number of countries, and other nations may have policies deterring paying foreign cloud providers. The challenges are even more acute in countries without either robust research computing infrastructure or the means for researchers to pay for computing resources.

To address the need for best practices and appropriate infrastructure, we combined the expertise of three projects—the Vertebrate Genomes Project (VGP), the European Reference Genome Atlas (ERGA), and the Galaxy Project—to create a global platform for eukaryotic genome assembly. The VGP is a collaborative effort to generate high-quality reference genomes for all vertebrate species (>70,000)^7^. ERGA is a pan-European scientific response to current threats to biodiversity, which aims to generate high-quality reference genomes for all ∼200,000 European eukaryotes, where about one-fifth of the assessed Red List species are at risk of extinction^2^. Galaxy^8^ is an open web-based scientific workbench made available through public computational infrastructure that has been delivering a robust and free analysis environment for the past 16 years. The Galaxy Project allows users to execute complex workflows on thousands of datasets and terabytes of data via either a graphical user interface or programmatically. Major global Galaxy instances in the United States (US), European Union (EU), and Australia provide access to powerful public computational resources. This unique combination of knowhow and infrastructure lays the foundation for democratizing genome assembly through universally and freely accessible assembly workflows.

The VGP has a long history of commitment to developing tools and workflows for high-quality reference genome assembly, including broadly employed assembly pipelines^1, 7^. The previously published VGP pipeline (v1.7) used Pacific Biosciences (PacBio) Continuous Long Reads (CLR) to generate a set of phased primary and alternate pseudo-haplotype contigs (or fully phased contigs when parental data were available)^9^ along with 10X Genomics (10X) linked-reads, Bionano Genomics (Bionano) optical maps, and chromatin conformation (Hi-C), capturing long-range information to join contigs into larger chromosome-level scaffolds. Given the lower accuracy of CLR reads compared to other sequencing data types, these scaffolds were then polished for base call errors with a combination of long and short reads.

While VGP has already assembled over 100 high-quality vertebrate genomes through this pipeline, recent radical changes in sequencing technologies—including the advent of PacBio high fidelity (HiFi) reads^10^, the discontinuation of 10X linked reads, as well as the parallel evolution of genome assembly algorithms—now call for a major update and enhancement of the pipeline. Particularly, PacBio HiFi reads constitute a paradigm shift over previous PacBio CLR^10^, as the higher accuracy allows assembly of long, highly identical repetitive regions^11^. With a combination of Oxford Nanopore (ONT) ultra-long reads, HiFi reads enabled the completion of the first telomere-to-telomere (T2T) human genome^12, 13^. HiFi reads are circular consensus reads (CCS) which are nearly error-free except for simple sequence repeats: this accelerates the assembly process, ultimately generating genome assemblies of the highest quality with minimal resources. This has been demonstrated in the human genome reference^12^, and evidence is rapidly accumulating that the approach successfully applies to many other species, including those with even larger and more complex genomes^14, 15^. By exploiting these sequencing advances and the latest computational improvements in genome assembly, we developed a new version (2.1) of the VGP assembly workflow. Furthermore, we deployed the workflow on open and global Galaxy infrastructure to make it freely accessible to researchers worldwide. Here we present this new version of the VGP assembly pipeline in Galaxy and demonstrate its performance by assembling and analyzing the genomes of 51 species. These species were assembled using automatic workflows in a process at least an order of magnitude faster than past efforts that lacked such workflows.

## Results

### A versatile collection of workflows for genome assembly

The VGP-Galaxy pipeline 2.1 is optimized for the assembly of PacBio HiFi reads along with supplemental scaffolding and phasing data. It is organized into ten Galaxy workflows (Fig. 1a), which can be launched with a few clicks in the Galaxy web portal. The logic of the pipeline is to progressively refine and complement the initial assembly graph to avoid the loss of information resulting from collapsing the graph to linear sequences (Fig. 1b). The first stage of the pipeline is the generation of *k*-mer profiles of the raw reads to estimate genome size, heterozygosity, repetitiveness, and error rate necessary for parameterizing downstream workflows. The generation of *k*-mer counts can be done from HiFi data only (Workflow 1) or include data from parental reads for trio-based phasing (Workflow 2; trio is a combination of paternal sequencing data with that from an offspring that is being assembled). The second stage is the phased contig assembly. In addition to using only HiFi reads (Workflow 3), the contig building (contiging) step can leverage Hi-C (Workflow 4) or parental read data (Workflow 5) to produce fully-phased haplotypes (hap1/hap2 or parental/maternal assigned haplotypes), using hifiasm^16^. The contiging workflows also produce a number of critical quality control (QC) metrics such as *k-*mer multiplicity profiles^7^. Inspection of these profiles provides information to decide whether the third stage—purging of false duplication— is required. Purging (Workflow 6), using purge_dups^17^ identifies and resolves haplotype-specific assembly segments incorrectly labeled as primary contigs, as well as heterozygous contig overlaps. This increases continuity and the quality of the final assembly^17^. The purging stage is generally unnecessary for trio data for which reliable haplotype resolution is performed using *k-*mer profiles obtained from parental reads. The fourth stage, scaffolding, produces chromosome-level scaffolds using information provided by Bionano (Workflow 7), with Bionano Solve and/or Hi-C (Workflow 8) data and SALSA or YaHS scaffolding algorithms. A final stage of decontamination (Workflow 9) removes exogenous sequences (*e.g.*, viral and bacterial sequences) from the scaffolded assembly. Additionally, a dedicated workflow (Workflow 0) is available for mitochondrial genome assembly.

**Figure 1.**
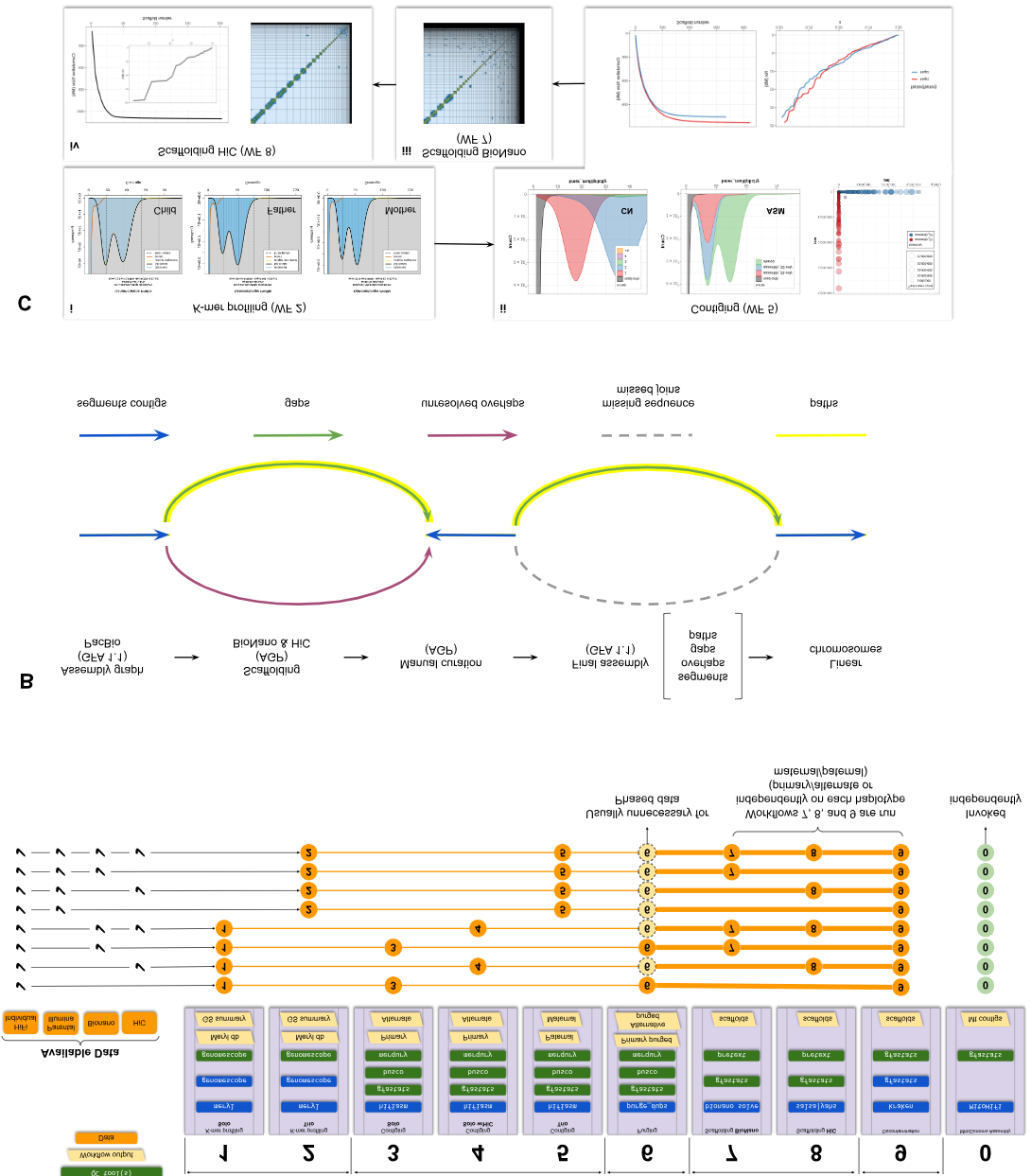
Assembly workflows, analysis trajectories, and quality metrics. **A**. Eight analysis trajectories are possible depending on the combination of input data. Decision on invocation of workflow 6 is based on the analysis of QC output of workflows 3, 4, or 5 (see Supplemental data for full explanation). Thicker lines connecting workflows 7, 8, and 9 represent the fact that these workflows are invoked separately for each phased assembly (once for maternal and once for paternal). **B**. Schematic of assembly graph propagation through the assembly pipeline using the GFA format. Initially, PacBio contigs are generated in GFA format. The scaffolding information from Bionano and/or Hi-C is added to the graph through AGP intermediates ^58^. Manual curation can be integrated in the graph using AGP files. The final assembly is a collection of segments and unresolved overlaps from the original graph, with gaps and paths representing the scaffolding information. Paths can be converted to linear FASTA sequences for downstream analyses. During the scaffolding process gaps (green arrows) are added as jump (J) lines between the segments (blue arrows) to the GFA. This allows the information on unresolved overlaps (purple arrows) to be maintained while missed joins (dashed arrows) inferred from the scaffolding information are added to the graph. The final linear sequences are represented as paths in the graph (yellow highlight). **C**. QC generated by the pipeline. Zebra finch trio data supplemented with Bionano and Hi-C data is shown. i. GenomeScope profiles using 21-mers on child HiFi data and parental Illumina data. ii. Merqury profiles (upper three graphs) and gfastats continuity metrics (lower two graphs) following contiging. The CN plot shows the number of copies of k-mers in both assemblies. Single copy k-mers correspond to heterozygous regions, and two copies to homozygous regions. This plot provides information about the potential need to purge the assembly if it shows the presence of k-mers in three or more copies. The ASM plot shows that the paternal and maternal assemblies share k-mers at ∼40× range that corresponds to the diploid coverage (also indicated by the rightmost peak in the child GenomeScope profile in panel). This is consistent with the expectation that phased assemblies will split the homozygous regions of the genome between the two haplotypes. Accordingly, heterozygous content in the genome is split mostly evenly between the two haplotypes, shown by the similar-sized assembly-specific peaks at ∼20× (the maternal peak is slightly bigger due to the presence of the Z chromosome, which is larger than the W in the paternal assembly). The assembly blob plot (rightmost graph in the top row of subpanel ii indicates excellent stratification of hapmers (k-mers specific to a particular haplotype) across maternal (red) and paternal (blue) assemblies. iii. Pretext map of maternal assembly after Hi-C scaffolding shows increase in continuity compared with the panel. iv. General statistics and Pretext map of the maternal scaffolding after a second scaffolding step using Hi-C data. The map shows improvements compared to the previous scaffolding step, with fewer scaffolds and less physical proximity between scaffolds (non-diagonal Hi-C signal).

For high quality genome assemblies, the VGP recommends at least 30× of each HiFi and Hi-C per haplotype (i.e. 60× coverage for diploid genomes).

### Validation in zebra finch

The zebra finch (*Taeniopygia guttata*) genome has been the focus of multiple in-depth analyses by the VGP^18, 19^. Numerous datatypes and existing benchmark assemblies are available for the individual bTaeGut2 (heterogametic, ZW), along with trio parental sequence data, making it an ideal test case for our workflows. We performed three types of assemblies using different combinations of data (**Supp. Table 1**): Solo assembly (Workflows 1, 3, 6, and 9) utilizes PacBio HiFi data for the single individual; Hi-C assembly (Workflows 1, 4, 8, and 9) adds Hi-C data for improved scaffolding; and lastly, Trio assembly (Workflows 2, 5, 8, and 9) adds parental (♂bTaeGut3 (homogametic, ZZ) and ♀bTaeGut4 (ZW)) Illumina data for haplotype phasing. Fig. 1c shows key quality control (QC) measures for trio assembly as automatically produced by the pipeline.

Overall, we found the HiFi data applied to different contig and scaffolding paradigms on Galaxy produced excellent assemblies directly from the workflows, with high contiguity, completeness, and accuracy. These draft assemblies were then manually curated to evaluate and resolve residual structural errors, remove contaminants, and assign sequences to chromosomes (**Supp. Material 1**). The main issues detected were manually fixed, including: (1) removal of false duplications in the Solo primary assembly due to incomplete purging; (2) manual rebinning of partially misbinned sex chromosomes and microchromosomes in the Hi-C-phased assemblies due to incorrect phasing of sex chromosomes in pseudoautosomal regions; and (3) reassigning some paternal-specific sequences that were mis-assigned to the maternal haplotype in the Trio assemblies, apparent in the *k*-mer spectra profiles (Fig. 2A and Fig. 2B). We identified the cause as contigs incorrectly assigned by Hifiasm^14^ based on too few parental *k*-mers. To fix this, we re-ran Workflow 5 specifying the problematic reads as ambiguous. After rebinning, we estimated the *k*-mer duplications and genome completeness of both haplotypes and found that the *k*-mer duplications were higher before rebinning (0.9% paternal and 4.4% maternal) than after rebinning (paternal 0.8% and maternal 3.7%) (Fig. 2C). Moreover, the rebinned paternal assembly showed 1.2% higher *k*-mer completeness than the default, although a slight decrease of *k*-mer completeness (0.2%) was shown in the rebinned maternal assembly (hapmer-specific completeness reported in **Supp. Table 2**).

**Figure 2.**
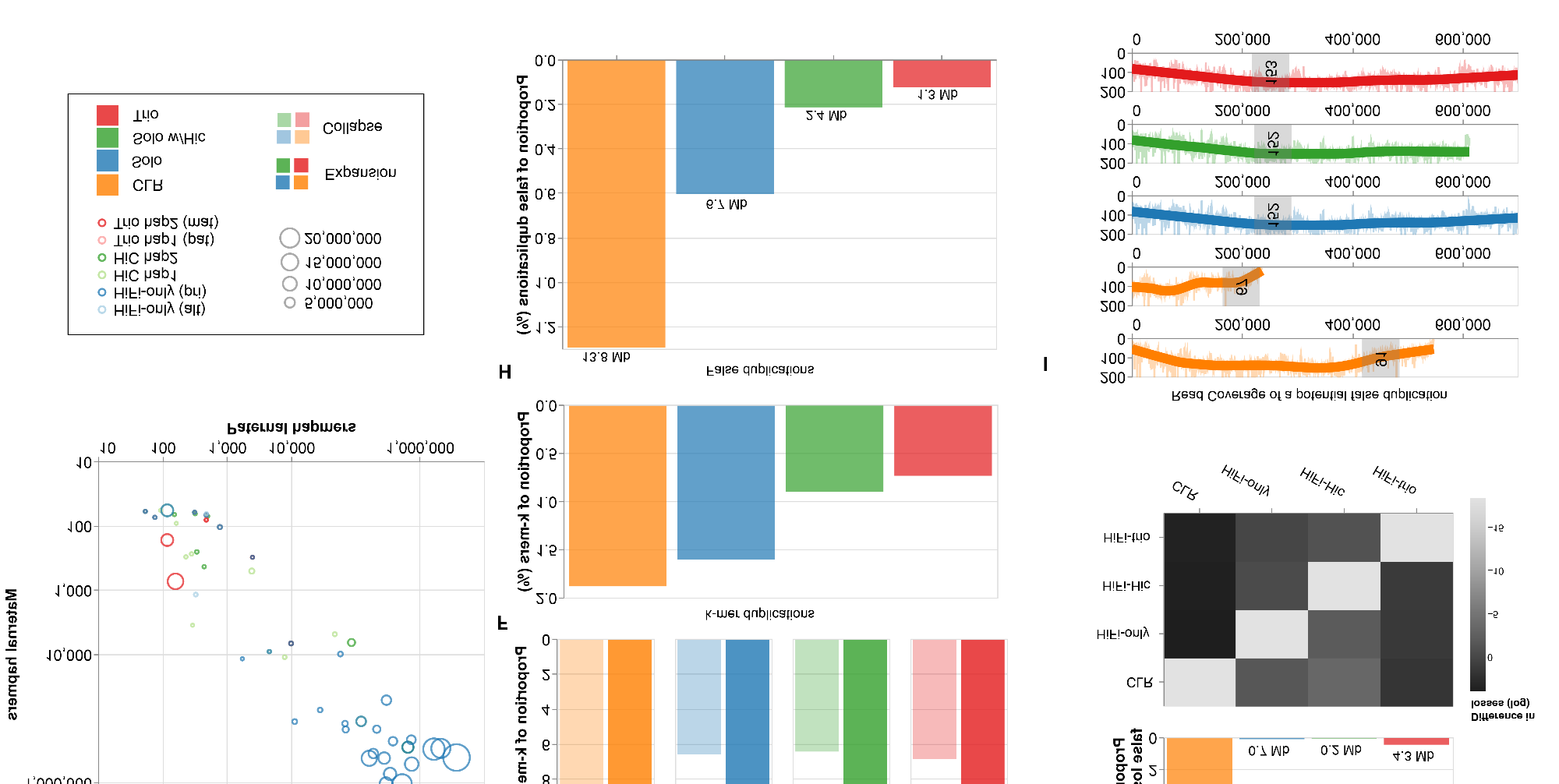
Extended evaluation of HiFi zebra finch assemblies using the three modes of the VGP pipeline in Galaxy: “Solo” (pseudohaplotype primary/alternate), “Solo w/Hi-C” (Hi-C-based phasing), and “Trio” (phasing using parental reads). **A**. shows 21-mers at 2-copy in the assembly with k-mer multiplicity between 50 and 90, a range that includes the expected diploid coverage region. **B**. shows k-mers that are present in only the reads, which suggests that the regions they represent (about 8 Mbp of sequence that ended up re-binned) are missing from the assembly, relative to the assembly in question. The purged Solo primary assembly has some missing regions, likely due to imperfect purging (k-mer multiplicity <5 excluded). **C**. Comparisons of k-mer duplication (red) and completeness (blue) between default and rebinned trio assemblies in males (left) and females (right). **D**. shows contigs from each assembly plotted according to how many parental hapmers are present in the contig, with contigs that were either fully phased (>95% of either parental k-mer) or lacking informative phasing information (i.e., less than 50 paternal and less than 50 maternal k-mers) excluded. The size of each bubble is proportional to the total k-mer size of the contig. Contigs along the diagonal have a mixed representation of hapmers from both parents, indicating intra-contig switch errors. Of these contigs, the Solo ones are typically larger and contain a higher amount of hapmers from both parents. **E**. Proportion of k-mer expansion and collapse in each diploid bTaeGut2 assembly. **F**. Proportion of k-mer duplication in the bTaeGut2 assemblies. We calculated k-mer duplications from the primary assemblies (CLR, Solo, Solo w/Hi-C) and paternal assembly (Trio) from phased diploid assemblies. **G**. Proportion and cumulated size (in Mb) of false losses of each assembly (above), and heat map of the size (in Mb) of false losses identified between the assemblies (below) in log scale. **H**. Proportion and cumulated size (in Mb) of false duplications of each assembly. **I**. A case of potential false gene gain in CLR assembly. Duplications of homologous sequences of partial ITSN1 gene was found in CLR assembly. Read depth coverage of contigs including the homologous sequences of ITSN1 gene in each bTaeGut2 assembly (highlighted in grey) is shown with a range from 0 to 200. The number in the gray highlighted region represents a mean depth coverage of ITSN1 homologous regions in each assembly.

To identify resolved, collapsed, or spurious gene duplications, we mapped HiFi reads to each HiFi assembly, and calculated read depth within RefSeq annotations^20^ (**Extended data Fig. 1**). We found 49 genes with multiple copies in the hap1 of the Hi-C phased assembly, and no more than one copy in each of the other assemblies. Of these genes, 41 have two copies in hap1 and no copies in the hap2 Hi-C assembly suggesting possible incorrect read phasing (**Extended data Fig. 2**). Notably aside from two genes in the hap2 Hi-C assembly, there are no other genes duplicated exclusively in one assembly. Genes with high copy numbers in each assembly were *ARL14EPL variant X2*, *PHF7*, and *VDAC3* with 19-43, 18-23, and 4-14 collapsed plus resolved duplications, respectively. Thus the same set of reads, assembled differently, gives a different number of genes and their copies when processed uniformly.

### Improvements of the HiFi-over the CLR-based assembly

The current reference of the zebra finch for the past several years, has been bTaeGutv1.4 (ZZ with added W), a CLR-based assembly generated by the VGP 1.6 pipeline^6^. We compared a CLR-based assembly to all of our HiFi-based assemblies generated in the VGP-Galaxy pipeline with a new individual, bTaeGutv2 (ZW). Based on *K\** statistics contrasting the *k*-mer frequencies of reads and assembly^21^, the CLR-based primary genome assembly contained the highest number of *k*-mer expansion and collapse errors (Fig. 2E) and the highest proportion of *k*-mer duplications (Fig. 2F). These analyses were confirmed based on analysis of whole genome alignments of CLR- and HiFi-based zebra finch assemblies and read depth calculations (see **Methods**), showing that the CLR-based assembly was more prone to apparent false duplications (Fig. 2G) and losses (Fig. 2H). The Trio HiFi assembly had the lowest amount of false duplications, whereas the Hi-C phased HiFi assembly had the lowest amount of false losses. The CLR-based assembly also contained more false gene gains and losses (**Extended data Fig. 3**). For example, the *ITSN1* gene, which is associated with autism-spectrum disorders^22^, was found to be partially duplicated in the CLR assembly (Fig. 2I), with 35 out of 39 coding exons found on two distinct contigs. Furthermore, for consensus quality evaluation (QV), we used *k*-mers from 10X short read datasets with Merqury^7^, and found that the HiFi Trio assemblies had consistently higher QV values (48.2 for maternal, 48.0 for paternal; **Supp. Table 3**) compared to CLR-based assembly (39.4 for the primary).

**Figure 3.**
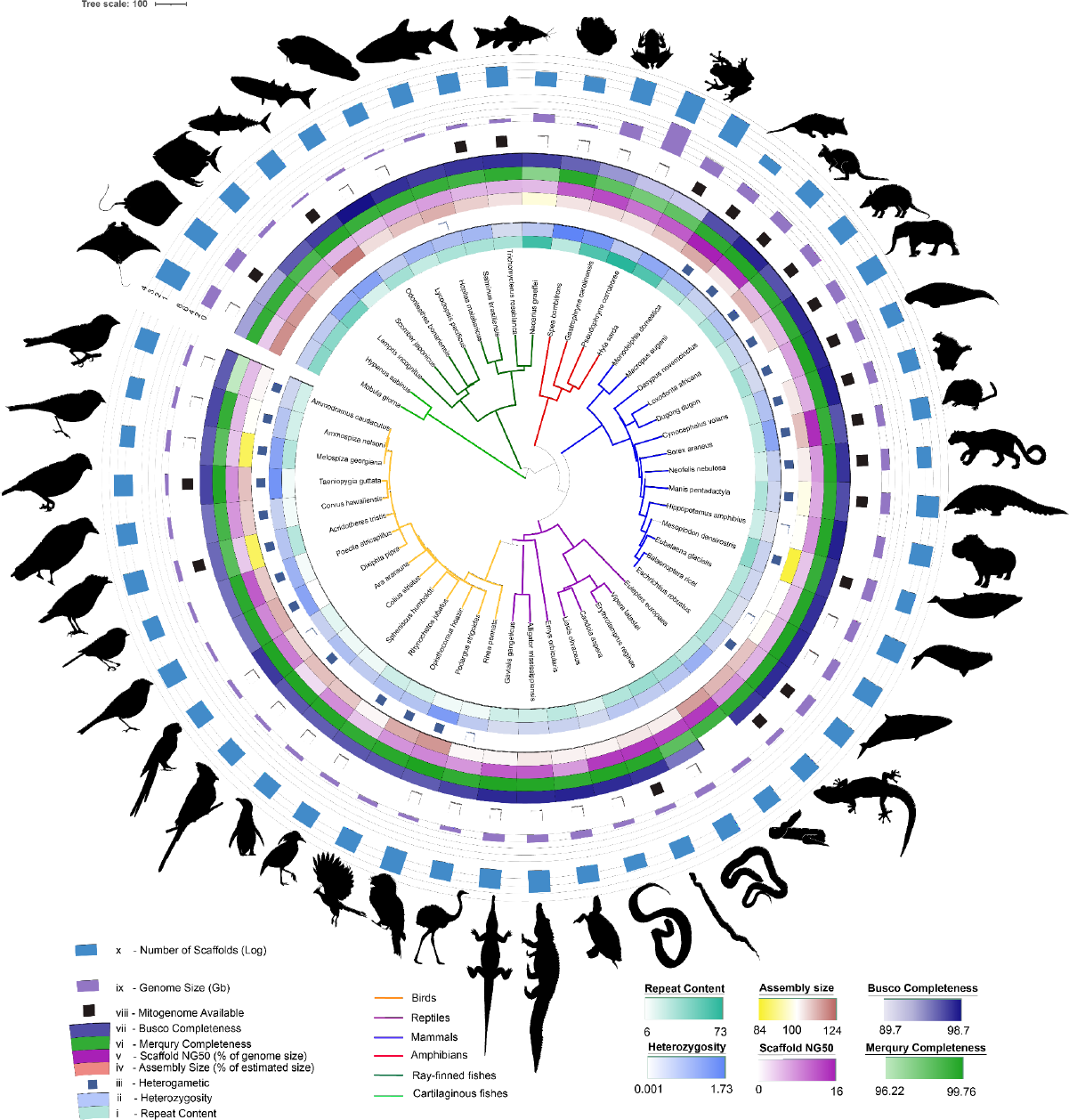
Phylogenetic tree and assembly statistics of genomes assembled using the VGP assembly pipeline in Galaxy. From the most inner circle to the most outer circle: (i) Repeat content, (ii) Heterozygosity, (iii) Individual with two identical sexual chromosomes (white) or two different sexual chromosomes (blue), (iv) Assembly size in percentage of the genome size estimated by genomescope, (v) Scaffold NG50 in % of estimated genome size, (vi) Merqury completeness of both haplotypes, (vii), BUSCO completeness: presence of orthologous genes present and complete compared to the set expected in vertebrates, (viii) Mitogenome assembled and available (black), (ix) Genome Size in Gb, with lines at 9, 2, 3, 4, 6, and 8 GB, (x) Number of Scaffolds in log scale with lines at 1 (10 scaffolds), 2 (100 scaffolds), 3 (1,000 scaffolds), and 4 (10,000 scaffolds).

### Hi-C data enables chromosome-level phasing

We tested whether trio-based phasing with parental sequencing data generates an assembly with more complete phasing than using Hi-C data^15^. Consistent with expectations, the trio assembly had the fewest contigs with mixed hapmer content (defined here as a contig having >10% paternal and >10% maternal hapmers), with two contigs from the maternal and two contigs from the paternal assemblies being mixed. The average size of these four contigs was 152,914 bp—an order of magnitude smaller than the average contig size of ∼1.5 Mbp. The Hi-C-phased assemblies had fewer contigs with mixed content (11 in hap1 and 7 in hap2), compared to the Solo haploid assemblies (26 and 1 in primary and alternate, respectively; Fig. 2D, **Extended data Fig. 5**). The size of contigs with mixed hapmer content was also an order of magnitude smaller in the Hi-C-phased assemblies (average size of mixed contigs being ∼0.87 Mbp compared to ∼1 and ∼2 Mbp for hap1 and hap2, respectively) compared to the Solo assemblies (average size of mixed contigs ∼6.8 Mbp compared to ∼1.6 Mbp for the primary). Since Hi-C data provide relative phasing information, but do not give actual haplotype-of-origin information, they cannot guarantee consistent phasing across separate chromosomes. Consequently, each haplotype in the Hi-C-phased assemblies has a mix of contigs from either the maternal or paternal haplotype, but the switch error rate^14^ within the Hi-C phased contigs is similar to that of the trio phased contigs (∼0.0016 and ∼0.0008 versus ∼0.0018 and ∼0.0007 for hap1 and hap2 for Hi-C and Trio, respectively; **Supp. Table 3**). After scaffolding each set of phased contigs separately with Hi-C and Bionano data, we observe that most of the scaffolds do not contain mixed hapmer content. This indicates that the Hi-C phasing from the start succeeded in properly binning whole chromosomes (**Extended data Fig. 5**), confirming what was previously reported for low-heterozygosity human data^15^. Due to its higher level of phasing and contiguity, we curated the Trio assembly to establish a new reference genome for zebra finch.

### Bionano Scaffolding marginally improves assembly quality when using Hi-C and YaHS

To evaluate the best scaffolding strategies using the current tools, we applied different scaffolding combinations on the HiFi generated contigs of the zebra finch we tested combinations of Hi-C scaffolding tools (SALSA2^23^ and YaHS^24^) with and without a Bionano optical mapping scaffolding tool (Bionano Solve tool), evaluating the number of gaps and scaffolds, as well as NG50 and auN^25^ (**Extended data Fig. 6, Supp. Table 4**). The combination of Bionano Solve and SALSA2 yielded a higher assembly contiguity than scaffolding with Bionano Solve or SALSA2 independently. However, YaHS performed similarly with and without Bionano information (auN values of 739.7 Mbp versus 732.5 Mbp for the wallaby primary assembly), and overall better than SALSA2. To test whether this result was influenced by species or sample characteristics, we performed the same test on two additional vertebrate species for which we had Bionano data: the Tammar wallaby (*Macropus eugenii*), a small marsupial macropod native to Southern and Western Australia; and the Snub-nosed viper (*Vipera latastei*), a threatened viperid snake species living in the Iberian Peninsula and Northern Maghreb^26^. The wallaby is of particular interest for scaffold-level assembly due to the large size of its chromosomes (∼100-730 Mbp in marsupials versus 50-250 Mbp in humans^27^). Scaffolding results in different taxonomy groups and genome sizes were comparable, overall leading to chromosome level assemblies (**Extended data Fig. 6, Supp. Table 4**). Our comparison also shows that when scaffolding Hifiasm generated contigs, the use of Bionano data only provides a marginal improvement when used with Hi-C YaHS scaffolding.

### Using workflows in production: assembly of 51 reference genomes

As part of VGP and ERGA reference genome assembly production projects, we used the pipeline to assemble 51 genomes: 14 mammals, 15 birds, 8 reptiles, 4 amphibians, and 10 fish (Fig. 3, **Supp. Table 5**). For all the assemblies, HiFi and Hi-C data were generated. The Hifiasm module was used for contiging and YaHS or SALSA modules were applied for scaffolding. Hi-C or parental short reads were used for haplotype phasing. These genomes were found to have a wide range of sizes (590 Mbp–8.5Gbp), repeat content (6%–73%), and heterozygosity (0.001%–1.73%). One of the species, the royal ground snake (*Erythrolamprus reginae*), was found during curation to be triploid, and has heterozygosity ranging from 0.422% to 3.74% depending on haplotype configuration (**Supp. Table 5**). To evaluate the overall quality of assembled genomes, we looked at gene- and *k-*mer-based completeness measures. On average, the assemblies showed ∼96% gene and ∼99% *k-*mer completeness with ∼1.93% of genes appearing as duplicated. The ratio of assembly lengths to the genome sizes estimated from *k*-mer profiling of the HiFi reads ranged between 0.84 and 1.24, with fishes exhibiting >1 ratios and birds <1. When we performed a similar analysis using Illumina sequencing of the same individuals for two species exhibiting HiFi-based high ratios (∼1.21 and 1.16 for *Scomber japonicus* and *Podargus strigoides,* respectively), the results were much closer to the observed assembly size (1.03 and 1.06, respectively). These findings suggest that some signal in the PacBio HiFi reads results in lineage species differences in *k*-mer estimated genome size and that this signal does not appear to be present in the Illumina k-mers estimate for genome size.

In agreement with the zebra finch results described above, the 51 HiFi-based assemblies overall had higher contiguity as a function of estimated genome size, repeat content, and heterozygosity when compared against 19 previous CLR-based genomes of other but comparable species produced with the first version 1.6 of the VGP-Galaxy pipeline (Fig. 4). Importantly, we observed that the HiFi-based assembly sizes were less impacted by heterozygosity and tended to be closer to the estimated genome size (Fig. 4D). Similarly to zebra finch, addition of Hi-C data improved haplotype resolution in other species. This was particularly striking for large, repeat-rich genomes such as Eastern narrow-mouthed toad (*Gastrophryne carolinensis*), where Hi-C phasing allowed full haplotype resolution without apparent false duplications (**Extended data Fig. 7A**). Due to the improvement in phasing brought by HiC-integration, all but one of the Hi-C-phased assemblies did not require running the purge_dups workflow according to the k-mer QC. For that reason, when parental data is not available, Hi-C phasing is the current standard approach in the VGP-Galaxy pipeline.

**Figure 4.**
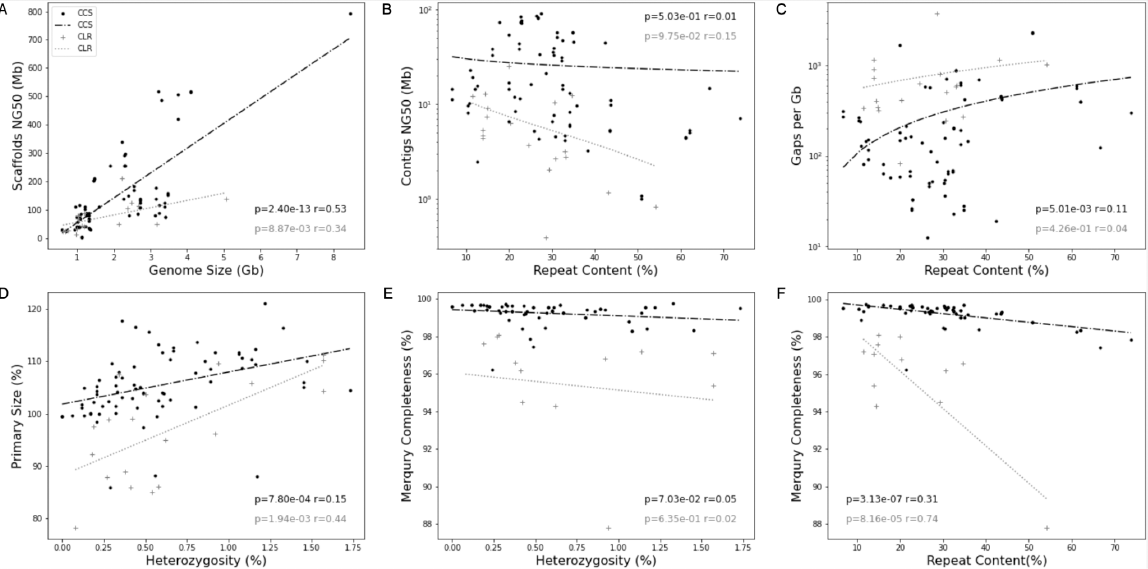
Comparison of assembly statistics between Sequencing technologies and scaffolding modes. Panels **A** to **F** compare assembly statistics between HiFi technology in black, used in this study, and CLR technology in grey, used in the previous version of the VGP assembly pipeline. Each dot represents an assembled species (Primary or Hap1), and the lines represent the linear regression for each technology. **A**. Scaffold NG50 in Mb in relation to Genome size in Gb. **B**. Contigs NG50 in relation to Repeat content, **C**. Gaps per GB in relation with Repeat content, **D**. Size of the Primary assembly in percent of the estimated genome size in relation with the heterozygosity rate of the genome, **E**. Genome completeness estimated by Merqury (both haplotypes together) in relation with the heterozygosity rate, F. Genome completeness in relation with repeat content.

### Contaminant analysis

The VGP decontamination pipeline is a new, automatic pipeline that we developed to remove exogenous sequences from assemblies. It was built and initially validated using 19 assemblies, which were chosen because they were decontaminated with the previous standard protocol (a combination of manual and automated screenings). The presence of contaminants or mitochondrial sequences was known and assumed to be ground truth (Fig. 5, **Supp. Table 6**). Contaminants represented a negligible fraction (<0.19%) of the assembled sequences (Fig. 5a, right side) and the contaminated scaffolds were generally small, with the largest being 0.15 Mbp, and the median number of bases removed was 61 kbp. Considering only assemblies with misclassified scaffolds (*n* = 5), where misclassification means the pipeline classification (assembly, contaminant or mitochondrial sequence) does not match the ground truth classification, the median size of false negative and false positive bases was 7.2 kbp (0.00051%) and 12.8 kbp (0.0012%), respectively (**Supp. Material 2**). After validation, the decontamination pipeline was used on 32 assemblies among the 51 described in this paper (**Supp. Table 7**). Of the 32, three assemblies contained foreign contaminants and five contained mitochondrial sequence. Kraken2^28^ uses a lowest common ancestor approach to classify contaminants, which can result in a high-level taxonomic classification. To better understand which foreign contaminants were identified, all contaminant sequences classified (from the 19 validation and additional three assemblies) were compared against the non-redundant nucleotide (nt) database, excluding one repetitive contaminant scaffold from *Erythrolamprus reginae* (rEryReg1) that was not masked by dustmasker and thus classified as a contaminant. The contaminants belonged to five different taxa, with *Escherichia* being the most abundant (Fig. 5b, **Supp. Table 8**, **Supp. Material 3**). Overall, these results show that our initial sequencing data had only trace amounts of contamination, and that the automated decontamination pipeline produces results as effective as the labor-intensive decontamination processes previously employed by the VGP.

**Figure 5.**
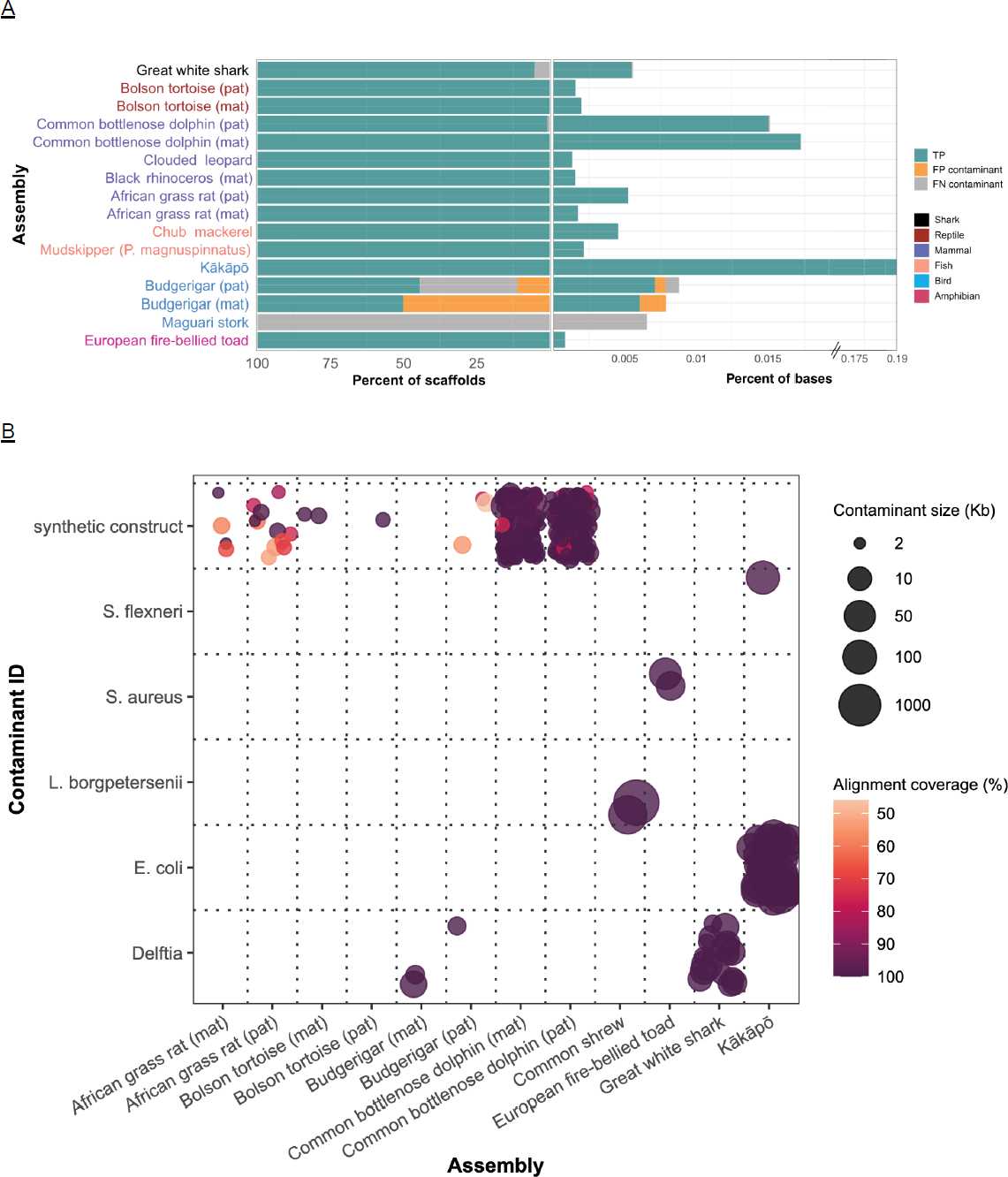
Decontamination pipeline. **A**. Comparison of foreign contaminants classified by VGP Decontamination Pipeline to the ground truth set. Percent of ground truth contaminant scaffolds that were true positive, false negative and false positive classifications, relative to percent of bases these scaffolds represent from the whole assembly. **B**. Species of contaminants identified by decontamination. Blast results of the foreign contaminants including coverage from the alignment and the size of the contaminant sequence. 323 of the 325 contaminant sequences identified are represented here; the two excluded contaminants were identified as killer whale and are false classifications. Two hundred sixty five of the contaminant sequences are synthetic constructs. The points are jittered to capture density.

### Mitogenome assembly and evaluation

Assembling complete and accurate mitochondrial genomes requires a distinct approach compared to nuclear genomes^29^, but can strongly support species identification and other analyses of mitochondrial evolution. Therefore, we implemented MitoHiFi (v. 2.2 and v. 3)^30^ as part of the VGP-Galaxy pipeline v2.1 to validate species identification and provide mitochondrial references. We tested MitoHiFi on the 51 species that were used for nuclear assembly. We then used the Barcode of Life Data System (BOLD) ID Systems^31^ for species identification and MITOS2 to annotate the reference as a functional evaluation (**Supp. Material 3**). We assembled 25 mitogenomes successfully and validated by annotation checks (**Supp. Table 9**). Mammals had the highest success rate (11/14), while fishes, amphibians, reptiles and birds had lower fraction of mitochondrial assemblies (3/5, 1/4, 4/8, 4/15 respectively). These results are congruent to those previously noted for PacBio CLR data^29^ in that the availability of mitochondrial reads appears to be a function of tissue type from specific taxonomic groups, as well as long-read library cut-off insert sizes smaller than 20 kbp (**Supp. Table 9**). For example, bird blood contains much fewer mitochondria than muscle, and vertebrate mitochondrial genome size is on average about 16 kbp.

Several assemblies also contained well-supported mitochondrial gene duplications (particularly in the control region *OH*) or missing tRNA genes (**Supp. Table 9**), as previously reported for other species^29^. These results allow us to conclude that, while HiFi reads provide an excellent approach to generate complete and accurate mitogenomes, scaling up to large volumes of samples (*e.g.,* for DNA barcoding purposes) will require careful consideration of experimental design in terms of sample selection and library preparation in order to retain mitochondrial reads.

### From new assemblies to biological insights

The phylogenetic breadth of genomes assembled here provides unique opportunities for studying evolution of their structure and function. While normally that would require generating multiple genome alignments—a complex and computationally expensive process—computation power underlying global Galaxy instances provides means for rapid analysis of dozens of new genomes that are in the process of generating RefSeq/ENSEMBL gene annotations. To demonstrate this, we generated a preliminary annotation of X-box protein 1 gene (*XBP1*) across our new assemblies, a transcription factor involved in regulation of protein folding of other proteins in regulatory regions of DNA, to trace down its evolution. We chose this gene for two reasons. First, its unique structure: *XBP1* mRNA is cleaved by a specific endonuclease, IRE1α, which removes a 26 bp spacer located within exon 4 (Fig. 6a) altering the phase of the reading frame downstream of the cleavage site^32^. This unique feature frequently renders this gene misannotated in genome assemblies. Second, the human genome contains an additional pseudogene copy of this locus. Our genome assemblies may shed light on when this duplication has occurred.

**Figure 6.**
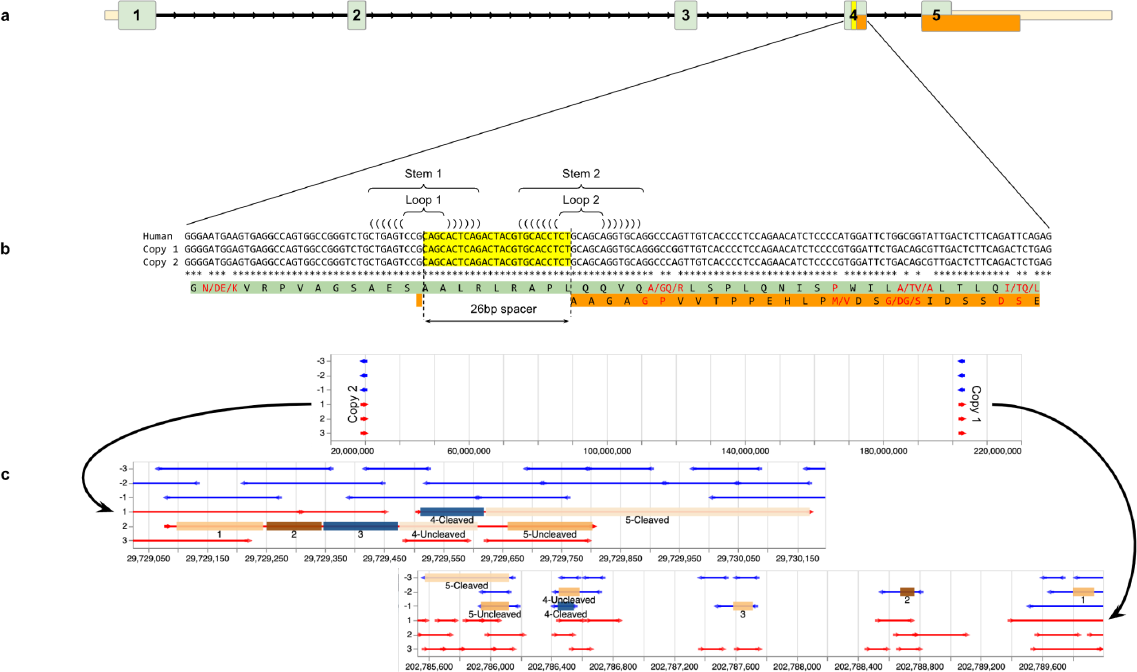
Duplication of *XBP-1* locus in *Cynocephalus volans* and comparison to the human locus. **a**. The structure of the human *XBP-1* gene. Exon 4 contains a 26 bp spacer (yellow) that is excised from mature mRNA by endonuclease IRE1α. When the transcript is not cleaved, the green reading frame is translated. When the spacer is removed the reading frame switches to orange downstream of the cleavage site. **b**. A nucleotide-level representation of exon 4 and corresponding translations for human and two flying lemur copies (Copy 1 and Copy 2). Single letter amino acid identifiers are centered at the second codon position. Red amino acids indicate sites with nucleotide changes. Two amino acids separated by “/” indicate amino acid replacement from one preceding slash to the trailing one: Q/R = change from Q to R. Single red amino acid indicates no change (synonymous substitution). Two stem-loop structures are critical secondary structure elements of the IRE1α cleavage site. **c**. Structure of two *XBP-1* loci in Cynocephalus volans. Top panel shows the relative position of the two copies within scaffold 2. Copy 1 retains exon/intron structure identical to that of the human gene. Copy 2 lacks introns completely. Arrows indicate all possible reading frames (STOP-to-STOP) in the vicinity of each exon. Red = + strand; Blue = - strand.

To identify this gene within assemblies described here (Fig. 3), we developed a workflow that finds reading frames with high similarity to exons of interest (see Methods). Interestingly, we found the gene experienced duplications in Philippine flying lemur (*Cynocephalus volans*), tammar wallaby (*Macropus eugenii*), and human (*Homo sapiens*; based on the analysis of the latest T2T-CHM13v2 assembly). In wallaby and human, only one copy is functional with the second copy being truncated and thus pseudogenized. However, in Philippine flying lemur there are likely to be two functional copies with distinct structures (Fig. 6b) located within the same linkage group (scaffold2) and separated by over 180 Mbp. Copy 1 appears to maintain the ancestral gene structure in comparison with the human orthologue, while Copy 2 lacks introns and appears to be a transposed copy (Fig. 6c). It likely arose via duplication followed by transposition of the reverse-transcribed uncleaved form of XBP-1 mRNA in a manner similar to the evolution of rodent insulin II gene^33, 34^. Both reading frames are present and appear as contiguous segments. Copy 1 and Copy 2 are ∼95% identical at the nucleotide level with two stem-loop structures required for IRE1α cleavage^35^ being absolutely conserved. The read depth in the vicinity of both copies appears to be uniform, ruling out assembly artifacts.

### Deployment, Scalability and Dissemination

Our goal was to both create and test a collection of assembly workflows and deploy them on public, freely available computational infrastructure to make genome assembly universally accessible to researchers worldwide. In parallel to using the public instance of Galaxy, we also successfully ran the whole pipeline on a local instance at the Rockefeller Vertebrate Genome Laboratory. The VGP Rockefeller University High Performance Computing (HPC) resources included 28 nodes each with ∼400 GB of RAM and 32 CPUs, as well as one node with ∼1.4 TB of RAM and 64 CPUs (**Supp. Table 10**). The pipeline instance maximizes time efficiency through Galaxy workflows’ parallelization of batch jobs as well as running non-dependent jobs concurrently to larger processes. The main run time bottlenecks were contig assembly with Hifiasm and mapping Hi-C reads with bwa-mem for YaHS scaffolding, which can take over a day of wall-clock time each depending on the genome characteristics. Additionally, Hifiasm has a longer runtime when using Hi-C phasing integration (Fig. 1; Workflow 4). Half of the evaluated species (all less than 3 Gbp estimated genome size) have their bottleneck runtimes total about a day or less, while the rest (all over 3 Gbp estimated genome size) ran in less than 5 days wall-clock time. After evaluation and fine-tuning we deployed workflows to the three global Galaxy instances (see https://galaxyproject.org/projects/vgp for information on how to access and use the workflows) in the US (https://usegalaxy.org), the EU (http://usegalaxy.eu), and Australia (https://usegalaxy.org.au). Each of the instances is supported by powerful “public cloud” infrastructure allowing multiple users to conduct numerous analyses simultaneously. For example, in the US, Galaxy is built on top of the Advanced Cyberinfrastructure Coordination Ecosystem: Services & Support (ACCESS-CI^36^) resources and has access to over 10 petabytes of storage and more than 50,000 cores, that can be leveraged to generate assemblies at scale. Moreover, Galaxy can be installed to run using local HPC systems, especially to better manage storage requirements and computing queues. Combined, these resource types provide a key advantage to reach the ambitious goals of the VGP, ERGA, and similar projects that require generating high-quality reference genomes at scale.

## Discussion

Producing high-quality reference genome assemblies for all 1.8 million named eukaryotic species is a daunting task. Accomplishing such an endeavor will require democratization of genome assembly so that it can be conducted by scientists regardless of their computational expertise or ability to access computational infrastructure. Large multinational efforts such as the VGP or ERGA excel in channeling methodological knowledge and technological advances into creation of best practices and have the capacity to refine these best practices in response to the continuous evolution of analytical methods and sequencing technologies. However, the best practices—ensembles of tools chained into workflows suitable for analysis of different types of input data—are not sufficient alone to truly democratize genome assembly. These workflows need to be readily accessible and supported by adequate computational resources. Here we address this challenge by combining analytical expertise of the VGP with the power of public computational infrastructure provided by the Galaxy Project in an attempt to make assembly universally accessible. We demonstrate that assemblies produced from HiFi reads surpass those produced with older CLR data in overall completeness and continuity. Our analyses underscore the value of Hi-C data for accurate haplotype resolution and increased continuity without the need for additional optical mapping information. Importantly, the work on optimization and streamlining of the underlying workflows allowed us to scale up the VGP assembly efforts.

Our approach to deployment of assembly workflows is designed to be useful across the full spectrum of user skill levels and analysis scenarios including: (1) biologists who need to assemble and analyze large genomes with a user-friendly system running best-practices approaches; (2) bioinformaticians and computational biologists who need to streamline, scale up, and automate assembly and alignment analyses using application programming interfaces (API) that can be run against public infrastructure; and (3) educators who will be able to use interactive tutorials developed within this project. The first group of users will benefit from using the graphical-user interface, while the second will be able to automate analyses by deploying them via command line scripts built using Galaxy’s Planemo software development kit^37^. Additionally, we compiled in-depth training tutorials that describe how to run the VGP assembly pipeline as well as in the interpretation of QC results (https://training.galaxyproject.org).

Future work will involve both improving the efficiency of the pipeline to minimize required resources, thereby increasing scalability, as well as continuously maintaining, updating, and upgrading the pipeline in response to improvements in the field. These changes will be needed to accommodate the dramatic increase in throughput of the PacBio Revio sequencer which can already sequence up to four mammalian genomes simultaneously with HiFi reads in one day for approximately $1,000 USD reagent cost per genome (for 30–35X coverage). Additionally, a particularly promising improvement is the integration of ONT data^38^ as an important complement to use in combination with PacBio for graph resolution^38^, and future work will include the integration of this technology in the assembly pipeline. Both long-read data types have strengths and weaknesses, and the combination of the two has shown promising results in the T2T assembly of a human genome^12^. However this approach needs to be tested and extended to additional taxa, including particularly challenging species and samples, such as those with genomes that are large, heterozygous, repetitive, and/or polyploid. Finally, a reference genome assembly often represents the first step for a study, and it is often followed by functional genomics analyses or comparisons to other species. As such, comprehensive genome annotation and comparative genomics workflows will be a focus of our future work.

## Online Methods

### Genome profiling (Workflows 1 and 2)

In Workflow 1 (for HiFi reads only) and Workflow 2 (For HiFi reads and parental illumina reads), *k*-mers are counted using Meryl^39^, and the *k*-mer profile is analyzed using GenomeScope2^40^. The input of this workflow is a collection of HiFi reads in FASTQ format^41^, the *k*-mer size, and ploidy.

### Phased genome assembly and duplicate purging (Workflows 3 - 6)

The logic of the pipeline (Fig. 1b) is to progressively refine and complement the initial assembly graph, taking advantage of the graph-based analysis and other functions introduced by gfastats^42^. The final product is a scaffolded assembly graph in the GFA1.2 format^43^. This approach constitutes a new conceptual framework in genome assembly, since it avoids the loss of information resulting from collapsing the assembly graph to linear sequences.

The initial contigging is done using Hifiasm^16^ (https://github.com/chhylp123/hifiasm). Hifiasm workflows generate *de novo* genome assemblies using three potential methods of phasing. The first mode, HiFi-only (Workflow 3), uses only the HiFi reads to build a haplotype-aware assembly. The primary and alternate assemblies are then purged (Workflow 6) using purge_dups^17^ (https://github.com/dfguan/purge_dups). The coverage histogram can be used to manually adjust cutoffs, if necessary. The second mode, Hifiasm-Hi-C (Workflow 4), uses Hi-C data to improve haplotype phasing. The third mode, Hifiasm-trio (Workflow 5), uses parental information to fully phase the haplotypes. All contigging workflows include quality control reports, using gfastats^42^ (https://github.com/vgl-hub/gfastats), BUSCO^44^ (https://github.com/WenchaoLin/BUSCO-Mod), and Merqury^7^ (https://github.com/marbl/merqury). On all test datasets Hifiasm was run with default parameters, except for the HiFi-only workflow where we turned off internal purging.

### Bionano scaffolding (Workflow 7)

The Bionano scaffolding workflow uses Bionano Solve^45^ for scaffolding and gfastats^46^ (https://github.com/ablab/quast) for quality control. On all test datasets, Bionano Solve was run with default parameters and without contig breaking (*i.e.*, excluding conflicting contigs during NGS-map conflicts).

### Hi-C scaffolding (Workflow 8)

The Hi-C scaffolding workflow can be used either on the primary contigs or on the scaffolds from Bionano scaffolding. Hi-C reads are aligned and prepared for scaffolding using the Arima mapping pipeline. Then SALSA2 ^23^ (https://github.com/marbl/SALSA) is used for scaffolding. Quality control is done with gfastats, BUSCO and PretextMap (https://github.com/wtsi-hpag/PretextMap) to visualize Hi-C contacts before and after scaffolding. On all test datasets SALSA2 was run with default parameters.

Alternatively, the workflow employs YaHS^24^ v1.2a.2 (https://github.com/c-zhou/yahs) and can also be applied to either contig or scaffold assemblies. Pre-scaffolding processing as well as quality control is the same as in SALSA2 Hi-C scaffolding.

### Decontamination pipeline (Workflow 9)

Masking is performed with dustmasker from NCBI Blast+ v.2.6.0 using dust level 40. Kraken2 v.2.1.1 identifies non-target contaminants with a confidence level of 0.3. Mitochondrial scaffolds are identified with blastn (v.2.6.0) and the NCBI Refseq mitochondrion database (release 212). To ensure a scaffold is entirely mitochondrial and not a NUMT, a custom tool (parse_mito_blast, v. 1.0.1) takes all alignments between unique scaffold-accession number pairs and calculates the total alignment coverage taking overlap into consideration. The threshold for classifying a scaffold as mitochondrial is 95% alignment coverage. A compiled list of contaminant and mitochondrial scaffolds is passed to the gfastats (v.1.2.2) --exclude-bed function to remove these scaffolds and remaining adaptors, and generate a new FASTA file. Eleven assemblies were used to test the pipeline during development. Validation was performed on an additional eight assemblies, including three that have no contamination or mitochondrial sequence.

### Mitogenome assembly and validation (Workflow 0)

To assemble mitogenomes from long reads, we implemented MitoHiFi in Galaxy^30^ (https://github.com/marcelauliano/MitoHiFi), which included creating a Docker image for MitoHiFi In addition to generating a functional mitogenome, this step also serves as a confirmation for taxon identification. All assemblies were run with default parameters. Validation consisted of confirming the species identification as well as functional validation. For species identification, we used the Barcode of Life Database (BOLD) identification engine to search the resulting mitogenome against all BOLD cytochrome c oxidase I (COI) barcode records, a standard method for species identification. To functionally validate the mitogenome, we used the MITOS annotation tool, available in MitoHiFI, to ensure that the mitogenome had all necessary and expected mitochondrial genes. MITOS annotation also was able to determine if there were any gene duplications, as seen with some tRNAs.

### Manual re-binning of trio assemblies

For the female zebra finch, since the problematic contigs from the maternal assembly (hap2) were absent from the paternal haplotype (hap1), we approached this by first trying to find the contigs that contain 2-copy *k*-mers which were present only in the maternal haplotype using meryl and meryl databases for each haplotype.

~~~
meryl print difference bTaeGut2_hap2_count bTaeGut2_hap1_count output \ bTaeGut2_hap2only
~~~

~~~
meryl equal-to 2 bTaeGut2_hap2_only output bTaeGut2_hap2_only_equalto2
~~~

~~~
meryl-lookup -existence -sequence bTaeGut2_trio.asm.dip.hap2.p_ctg.fa -mers \ bTaeGut2_hap2_only_equalto2 > bTaeGut2_hap2_only_equalto2.tsv
~~~

This TSV (tab-separated values) file contained all the contigs in the maternal haplotype, the contig’s size (in 21-mers), and how many *k*-mers were seen in both the contig and the database of 2-copy *k*-mers present only in the maternal haplotype. We used this data to calculate the percentage of each maternal contig that belonged to the database of problematic *k*-mers, and we focused on contigs with over 50% of their *k*-mers matching that database. We then used the original GFA output from Hifiasm to find the reads that were used to build these contigs. We created manual re-binning lists that assigned these reads as “ambiguous” and re-run Hifiasm with the trio data as well as these re-binning lists.

### Assembly comparisons

For the zebra finch assembly comparisons, we used the three primary assemblies (CLR, Solo, and Solo w/Hi-C mode) and one rebinned paternal assembly (Trio mode) made immediately after contigging and purging (https://genomeark.s3.amazonaws.com/index.html?prefix=species/Taeniopygia_guttata/bTaeGut2/). All assemblies were masked by repeatmasker (https://www.repeatmasker.org/; ^47^ with default engine and commands “`-species ’Taeniopygia guttata’ -xsmall -s -no_is -cutoff 255 -frag 20000`” before genome alignment. The reads produced for the assemblies by PacBio CLR, HiFi and 10X platforms were mapped to the all genome assemblies by Minimap2^48^ and EMA mapper^49^. We used parameter “-ax map-pb” for PacBio CLR read, and `“-ax map-hifi”` for HiFi read mapping using Minimap2. The paired-end 10X reads were mapped using the barcodes default options of EMA^49^. The reads without barcodes were mapped using BWA^50^ with parameters “-p -M -R ‘@RG\tID:rg1\tSM:sample1’” following guidelines in EMA. Intermediate BAM files produced in the read mapping step were merged by Sambamba^51^. Samtools^52^ was used to sort the BAM files and to calculate read coverages of each genomic position.

We calculated *k*-mer duplications of each assembly using a script “false_duplication.sh” in Merqury^7^ with optimum *k*-mer size of the zebra finch genome: 21. We calculated *k*-mer collapse and expansion with Merfin^53^ using the same *k*-mer size. To compare the *k*-mer collapses and expansions of diploid assemblies, we included the maternal or alternate zebra finch sequences with the paternal or primary sequences. For optimum K* calculation, we included “lookup_table” produced by GenomeScope2^40^ with the 10X reads. We included a current reference genome of zebra finch assembled from CLR reads (bTaeGut1.4; GCF_003957565.2) in this analysis. We estimated *k*-mer duplications and completeness of default-mode and rebinned trio assemblies using both paternal and maternal assemblies without purging with 10X reads using Merqury.

We identified false duplications and losses using whole genome alignment with estimation of number of paralogs in alignment blocks. Firstly, we aligned the three primary assemblies (CLR, Solo, and Solo w/Hi-C mode) and the one paternal assembly (Trio mode) of the zebra finch using the Cactus alignment tool^54^. Then we extracted homologous regions to a readable multiple alignment format using HAL^55^. Because all assemblies were made from the same sample, the number of paralogs of each assembly in each alignment block should be the same, and when not the same, they are false duplications or losses occurred in one of the assemblies. To every alignment block showing this discordance of the number of paralogs between the assemblies, we calculated the likelihood of each number of paralogs to model (i.e. how many paralogous sequences will be present in the alignment block) based on summed read-coverage of the PacBio CLR, HiFi, and 10X reads. The likelihood of each model was calculated as

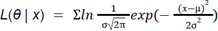

where *x* is the sum of mean depth of each homologous sequences in an alignment block from an assembly, and µ and σ are the parameters of depth distribution estimated from each number of paralogs models. To estimate the model parameter µ and σ, we calculated the mean and variance of normal distribution from depth coverages of genomic regions that there is no multi-copy *k*-mer for the model that the number of paralogs is zero, then simply multiplied the mean by integer for each model, e.g. multiplying the mean by 2 for 1-paralogs model. We supposed that the variance is the same for all models. The false duplications and losses of each assembly were identified when each assembly had more or less paralogs than a best model from the likelihood estimation in each alignment block. To remove the noise for the false duplications and losses, we filtered out false duplications on contigs where false duplications occupied <50% of the contig length, and far from terminals of the contig (>20kbp), and filtered out false losses under 1kbp length. To avoid false losses include haplotype differences we calculated *K*\* of *k*-mers in the region of candidate false losses after noise filtering mentioned above. We only included the candidates to false losses when a candidate has collapsed *k*-mers (K* >0) above 90% of the genomic sequences. Moreover, we estimated potential false gene gains and losses based on annotation data of bTaeGut2.trio (GCF_008822105.2) and the erroneous regions we identified in each assembly. We aligned the bTaeGut2.trio assembly and others together using Cactus ^54^, then, potential false gene gain or losses were identified when the false duplications and losses had homologous regions with any CDSs of bTaeGut2Trio annotation.

### Comparative gene analysis

For each genome we annotated all open reading frames (ORFs) ≥99 bp (defined as uninterrupted runs of sense codons bound by stops) using orfipy^56^. Next, we used amino acid translations of ORFs to create a Diamond^57^ database. We then queried the database using amino acid translations of human exons to identify the most likely location in assembled genomes and plotted this information using Galaxy/Jupyter integration to generate Fig 6. The Galaxy history with the step-by-step description of this process is available at https://gxy.io/GTN:T00174.

### Accessing the workflows

In addition to producing high-quality reference assembly as part of the Vertebrate Genome Project, the purpose of this work aims to democratize assembly for the entire community. This goal is achieved by making tools and workflows available on public instances where access to the computing resources is free. The analyses described here have been run on public Galaxy instances: usegalaxy.eu in Europe, with a dedicated assembly portal (https://assembly.usegalaxy.eu/), and https://usegalaxy.org in the United States. Some genomes have been assembled on a VGP Galaxy instance installed on lab servers, especially within the Vertebrate Genome Laboratory (VGL) at Rockefeller University. The workflows are accessible to everyone (https://galaxyproject.org/projects/vgp/), and the latest version is listed on the Galaxy project VGP page (**Supp Table 11B**). This page describes the workflows and their usage in detail along with educating users unfamiliar with genome assembly through a set of two training materials. A short version describes the pipeline superficially for a quick use of the workflows (**Supp. Table 11C**), and a long version goes in depth with every tool and parameters selected inside these workflows (**Supp. Table 11D**).

## Author contributions

D. L. built the assembly pipeline with support from G. F., L. A., C. G., B. G., A. O., H. C., M. D. S., B. D. P, A. R., M. V. D. B., and the VGP assembly working group. L. A., A. D., G. R. G., A. M. G., G. M. G., N. J., C. J., B. O., D. D. P., S. S., M. S., and T. T. generated one or several assemblies used in the analyses. B. J. K., K. R., and M. C validated the zebra finch assemblies. J. C. performed the manual curation on the zebra finch assembly. L. A. assembled and evaluated the mitochondrial genomes. N. B. established the decontamination pipeline and performed the contamination analyses. N. B. and M. P-F. compared the scaffolding strategies. A. N. performed the analyses on XBP1. C. G. and B. D. P. developed the training material with support from the user community. J. R. B., N. J., T. T., B. O., O. F., C.L., H. K., T. M-B, and R. M. W. generated the PacBio and Hi-C data. G. F., M. C. S., A. N., A. M. P., and E. D. J., conceived the study and drafted the manuscript. All authors contributed to the manuscript and approved it.

## Data Availability

These genomes were supported by collaborators of the VGP and ERGA, and quality control analyses reported here to test the VGP Galaxy pipeline does not release those that are under specific embargo policies for genome wide analyses (e.g. https://genome10k.ucsc.edu/data-use-policies/).

## Supporting information

Supp. table Summary

## Acknowledgements

We thank Yagoub Adam, Tyler Alioto, Jun Aruga, Sagane Dind, Diego Fuentes, Shilpa Garg, and Jèssica Gómez for contributing to the initial implementation during the ELIXIR Biohackathon 2021. We also thank Nate Jue for help testing and developing the pipeline tutorials, as well as Agostinho Antunes and Andrea Guarracino for their useful comments to the manuscript. We are thankful to Kathleen Horan and Melanie Couture for their work sourcing and coordinating the samples used for this paper. This work was supported in part by the Intramural Research Program of the National Human Genome Research Institute (NHGRI), National Institutes of Health (NIH), and Howard Hughes Medical Institute (HHMI). The authors are grateful to the broader Galaxy community for their support and software development efforts. This work is funded by NIH Grants U41 HG006620, U24 HG010263, U24 CA231877 and U01CA253481, along with NSF Grants 1661497, 1758800, and 2216612. This work was also supported in part by The Human Frontier Science Program (HFSP) RGP0025/2021, the Swiss National Science Foundation (SNSF) grants 202669 and 198691, the Swiss State Secretariat for Education, Research and Innovation (SERI) grant 22.00173 and Horizon Europe under the Biodiversity, Circular Economy and Environment program (REA.B.3, BGE 101059492). Usegalaxy.eu is supported by the German Federal Ministry of Education and Research grants 031L0101C and de.NBI-epi to B. G.

