## Extended data for "Scalable, accessible, and reproducible reference genome assembly and evaluation in Galaxy"

### Supplementary Material

#### 1. Manual curation

All variations of the bTaeGut2 assembly generated by the pipeline (*i.e.*, the solo, HiC-phased, and trio assemblies) have been manually curated to evaluate and resolve residual structural errors, remove contaminants, and assign sequences to chromosomes.

##### Solo assembly

Alignment to bTaeGut1.4.pri (GCA_003957565.4) showed that the primary bTaeGut2 assembly had the expected karyotype and that the smallest of the microchromosomes (31 - 37) were highly fragmented. Curation of the solo assembly resulted in the manual correction of scaffolds resulting in 31 scaffold breaks, 90 joins and approximately 11.7Mb of sequence removed as haplotypic duplication. The curated genome yielded 39 autosomes plus Z and W, with 98.55% of the sequence being assigned to chromosomes. Retrospective alignment of this assembly to the Hi-C phased assembly from the same sample revealed 9Mb of sequence missing from the W chromosome, this missing sequence appears to have been removed by the purging process.

##### HiC-phased assemblies

The HiC-phased assemblies were curated as individual haplotypes. Alignment of both haplotypes to the Solo and Trio assemblies of bTaeGut2 revealed poor haplotype separation particularly in relation to the sex chromosomes and the smallest of the microchromosomes. In part this may account for the difference in size between the larger Hap1 1.16Gb assembly versus the 1.06Gb Hap2 assembly. Incomplete representations of Z and W were found in both haplotypes requiring manual redistribution totalling approximately 44.7Mb of sequence in order to obtain single, complete representations for Z and W. Regarding microchromosomes 29 through 37 it was found that Hap1 contained virtually complete, albeit fragmented, representations for these plus over 60 duplicated sequences amounting to approximately 6.28Mb missing from the microchromosomes in the Hap2 assembly. It was also noted that a handful of sequences appeared to be represented only once between both data sets indicating incomplete phasing over these regions. Artificial triplication as an artefact of the phasing process led to the complete removal of 2.8Mb of sequence from the assemblies.

For each of the HIC-phased assemblies manual curation ultimately resulted in the expected karyotype of 39 autosomes plus Z or W. Manual interventions for Hap 1 amounted to 37 scaffold breaks, 159 joins and a total of 9.99Mb of haplotig removals for transfer into Hap2 with 95.99% of the sequence assigned to chromosomes. For Hap2 there were 30 scaffold breaks, 82 joins and a total of 23.7Mb of haplotig removals for transfer into Hap1, 97.93% of the sequence was assigned to chromosomes.

##### Trio assemblies

The Trio Maternal and Paternal assemblies were curated individually. The assemblies exhibited good separation of haplotypes with each assembly achieving the expected karyotype. As observed for other iterations of this assembly the smallest chromosomes; 30-37 were fragmented. For the Maternal assembly manual correction of the assembly resulted in 16 scaffold breaks, 88 joins and 0.3Mb of sequence removed as haplotypic duplication with 99.53% of the sequence assigned to chromosomes. For the Paternal there were 32 scaffold breaks, 193 joins and 0.6Mb removed as haplotypic duplication, 99.33% of the sequence was assigned to chromosomes.

#### 2. Assembly Decontamination

We introduced an automated decontamination pipeline to remove foreign DNA from viruses, bacteria and symbionts, as well as high copy mitochondrial sequence. Our validation assemblies included a total of 325 foreign DNA contaminants and 19 mitochondria. Only 3 of the 19 assemblies had no foreign DNA or mitochondria and were used to check for spurious results. Out of the 340 sequences that the decontamination pipeline identified as potential contaminants, 337 were true positive identifications (99.1%, 337/340, including mitochondria) and three were false positives based on our benchmark. Seven contaminants were missed (2.0%, 7/340) (Figure 5A). All 19 mitochondrial sequences were identified. Overall, the pipeline was able to exactly replicate the results of manual decontamination for 14/19 assemblies.

The five assemblies where our decontamination results differed from the benchmark include the primary assembly of the Great White shark, the paternal haplotype of the Bottlenose dolphin, the maternal and paternal haplotypes of the Budgerigar and the primary assembly of the Maguari stork (**Figure 6b**, right side). The largest incidence of misclassification was observed in the Maguari Stork, where there were two contaminant sequences (both *Neospora caninum*) in the benchmark and neither were identified, but the scaffolds only represent 0.0065% of the length of the genome (81.7Kb/1.25Gb). *N. caninum* and other parasites are likely not present in the Kraken2 database, and thus would not have been identified by the pipeline. In the case of the Budgerigar, two scaffolds in the maternal assembly and one scaffold in the paternal assembly were false positive classifications relative to the benchmark. The benchmark set is based on the prior VGP's established decontamination process, which involves manual and automated systems. While the results of this process are generally reliable, it is possible that true contaminants were not originally identified and thus included in the benchmark set. Based on a BLAST query against NCBI's nucleotide (nt) database, these three scaffolds are likely true contaminants with high similarity (>99% coverage) to *Delftia* species. The paternal assembly also contained three false negative classifications. These false negative scaffolds contain large segments of homopolymers and low-complexity repeats. These would have been masked in the assembly submitted to Kraken2, and it is possible that the remaining sequence was not long enough for reliable classification. When isolated, these non-repeat regions show high sequence similarity (>99%) to a synthetic construct from PacBio, which is a positive control sequence used for determining run success, so these are likely true contaminants that the pipeline did not identify. Interestingly, a BLAST search based on alignment coverage identified 150 contaminants from the Budgerigar, Bottlenose dolphin, Bolson tortoise and the African grass rat as almond (*P. dulcis*, accession AP021729). Further investigation revealed that this was caused by residual contamination from the PacBio internal control sequence (accession MG551957) in the published almond genome. In total, 265 contaminants were identified as the PacBio control sequences.

#### 3. Mitogenome assembly

MitoHiFi uses a reference mitochondrial assembly to select HiFi reads that map to the reference, which is suggestive of their putative mitochondrial origin (though nuclear mitochondrial DNA — NUMTs — and other nuclear material can be a confounding factor) [1]. The reference mitochondrion can come from the same species, or a closely related one if not available. MitoHiFi then filters the reads and uses hifiasm to assemble the mitochondrial genome, and further steps circularise and annotate it. It is also possible to use MitoHiFi on pre-assembled contigs. We tested MitoHiFi on the raw reads for the 51 species assembled for this paper (ST9). When MitoHiFi failed, it was usually due to a lack of true mitochondrial reads resulting in a failed assembly attempt; though there were cases when a poor quality mitochondrial assembly was generated from sequences likely belonging to NUMTs. NUMTs can be differentiated by usually being linear sequences longer than 16 kbp [1].

There were three cases where MitoHiFi failed to assemble a functional mitochondria, and further investigation yielded a proper mitogenome assembly: bMorBas2 (*Morus bassanus*), mMusLut2 (*Mustela lutreola*), mDasNov1 (*Dasypus novemcinctus*), and mCynVol1 (*Cynocephalus volans*). For bMorBas2, the problem was that *Morus* is the name of both the genus for gannets as well as mulberries, so the program had initially used a mulberry sequence as reference. For mDasNov1 and mMusLut2, the program selected a concatamer assembled from genuine mitochondrial reads, so this was fixed by removing the duplicate sequence. For mCynVol1, the MitoHiFi-produced assembly worked for BOLD [2] identification, but MITOS2 indicated internal stops in *COX1*, *COX2*, *ATP6*, *COB*, *NAD3*, and *NAD4*, and further the mitochondrial assembly selected was a linear contig that corresponded to a 33 kbp linear unitig. Inspection of the raw unitig graph generated by the hifiasm step in MitoHiFi indicated the presence of a 16,762 bp circular unitig, a more likely candidate for a mitogenome.

We tested MitoHiFi on 51 HiFi datasets. We then used BOLD ID Systems [2] for species identification based on a 648-bp region of the cytochrome c oxidase I (*COI*) gene and MITOS2's functional annotation[3] to evaluate the assembly quality with regard to missing genes or duplications. Resulting assemblies fell into three categories:

1. Complete and accurate assemblies: proper species identification in BOLD and no strong peculiarities reported by MITOS2 (25/51);
2. Further investigation (1/51);
3. No Assemblies: MitoHiFi failed to produce an assembly at all (25/51).

### Extended Data Figures

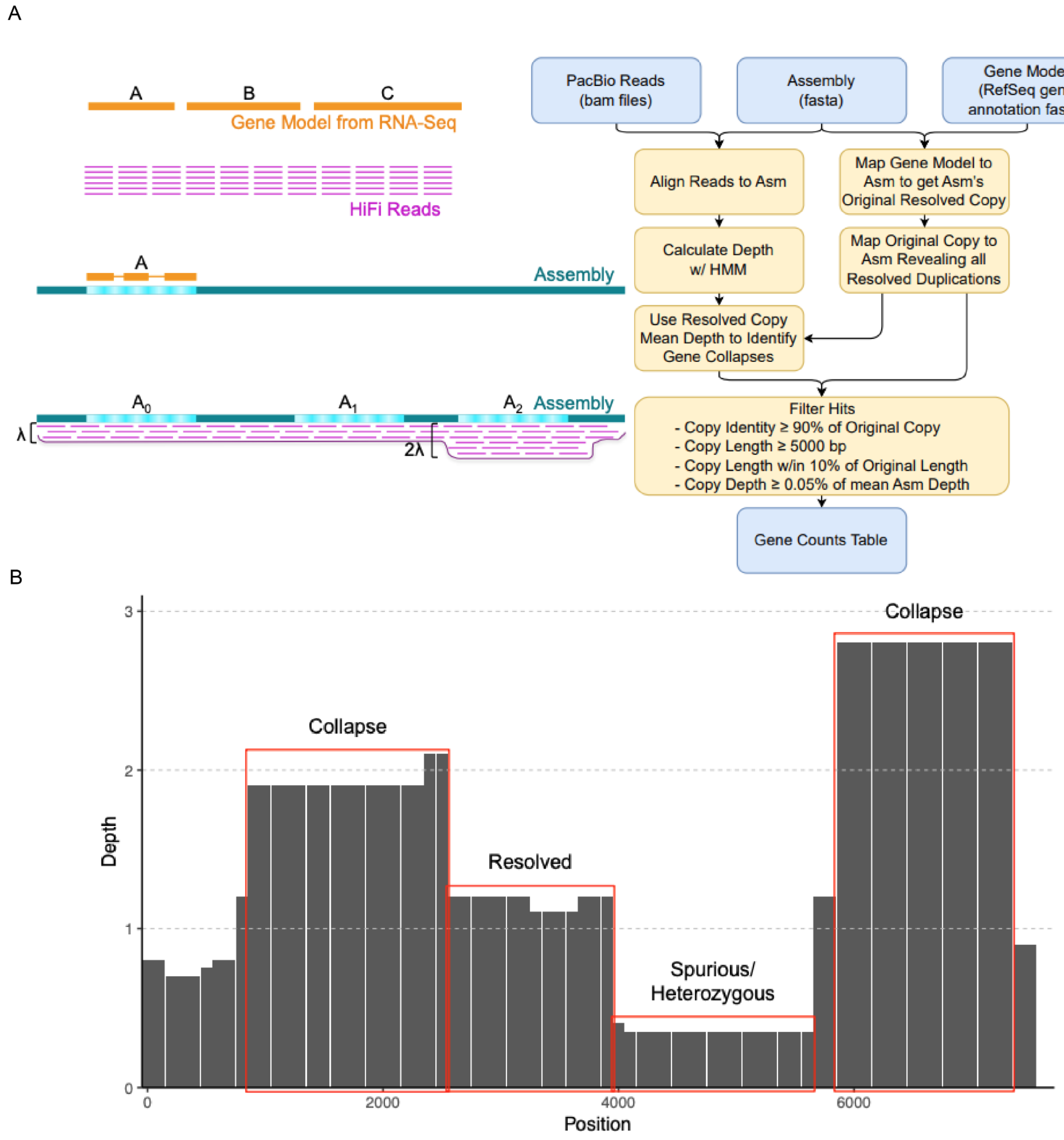


**Figure S1** | a) Method to obtain gene count tables of duplicated and collapsed genes. We aligned the PacBio HiFi reads to each assembly and used an hmm algorithm to calculate the read depth in sliding windows (Figure S1). We mapped the RefSeq gene Model to each assembly to get the assembly's original resolved copy. We then used the hmm depth to calculate the Mean Depth of resolved copies to identify gene collapses. We filtered the genes showing discordant depth to keep genes where the copy identity is above 90% of the original copy, the copy is longer than 5 kbp and within 10% of the original length, and the discordance in read depth is higher than 0.05% of the average assembly depth. b) Three types of regions are identified by the hmm algorithm: i) Resolved regions if the read depth is close to the assembly average depth, ii) Spurious duplications or heterozygous regions if the read depth is below the average, iii) Collapsed duplications if the read depths is twice the average or above.

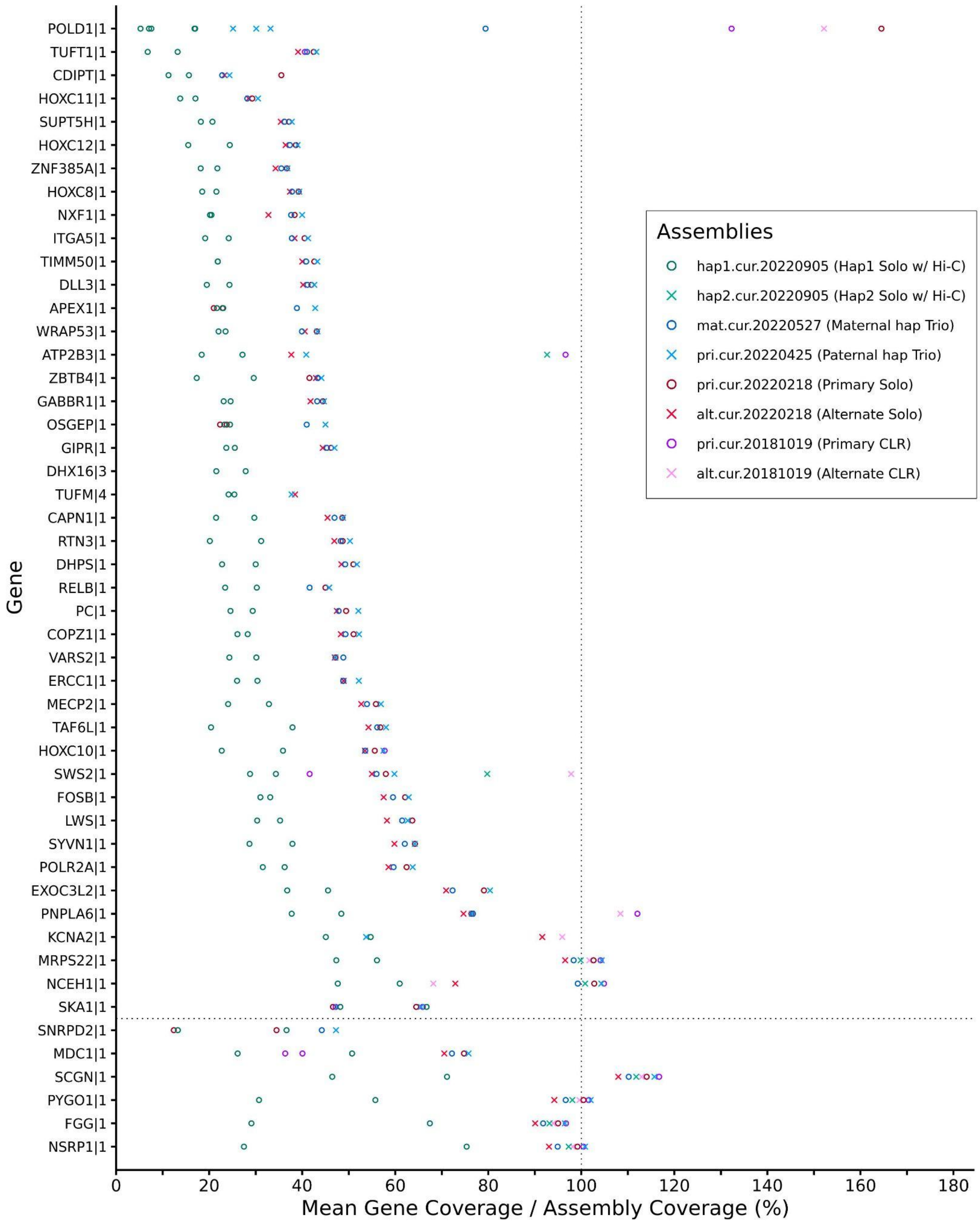


**Figure S2 |** Spurious Duplications in the Hi-C phased assembly of the zebra finch. Each data point of this graph represents a copy of the gene in the Y axis. The X axis represents the average read coverage relative to average coverage across the assembly. This graph shows that most of the spurious duplications occur in low coverage regions, and we can see that the two copies present in hap1 of the Hi-C assembly (green circles) have on average half the read coverage compared to the single copy in the other assemblies.

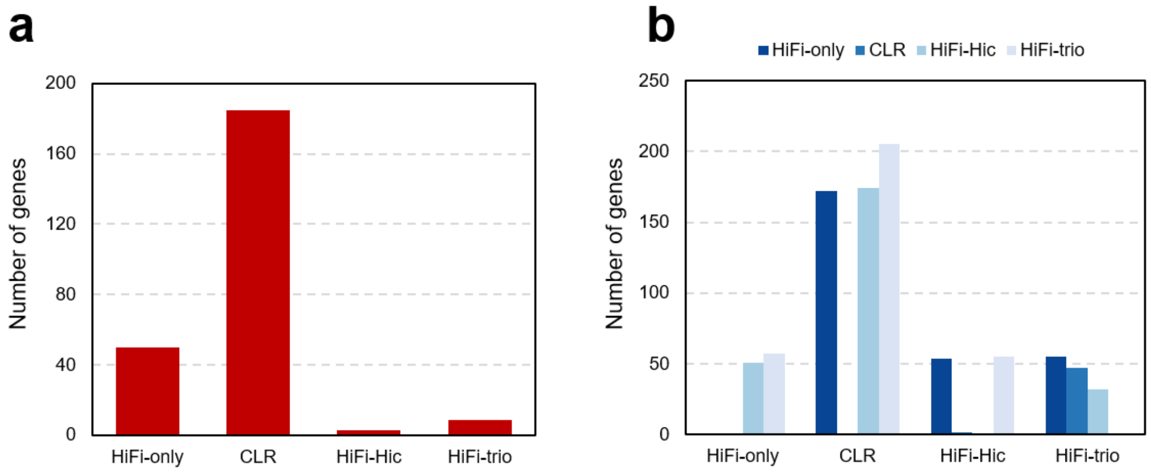


**Figure S3. a.** Number of potential false gene gains in each zebra finch (bTaeGut2) assembly. Potential false gene gains are calculated for individual genomes based on false duplication identified by read coverage of each assembly. **b.** Number of potential false gene losses in each bTaeGut2 assembly (X-axis) estimated from other bTaeGut2 assemblies (colours). False losses are estimated by comparing missing regions between two assemblies from whole genome alignment.

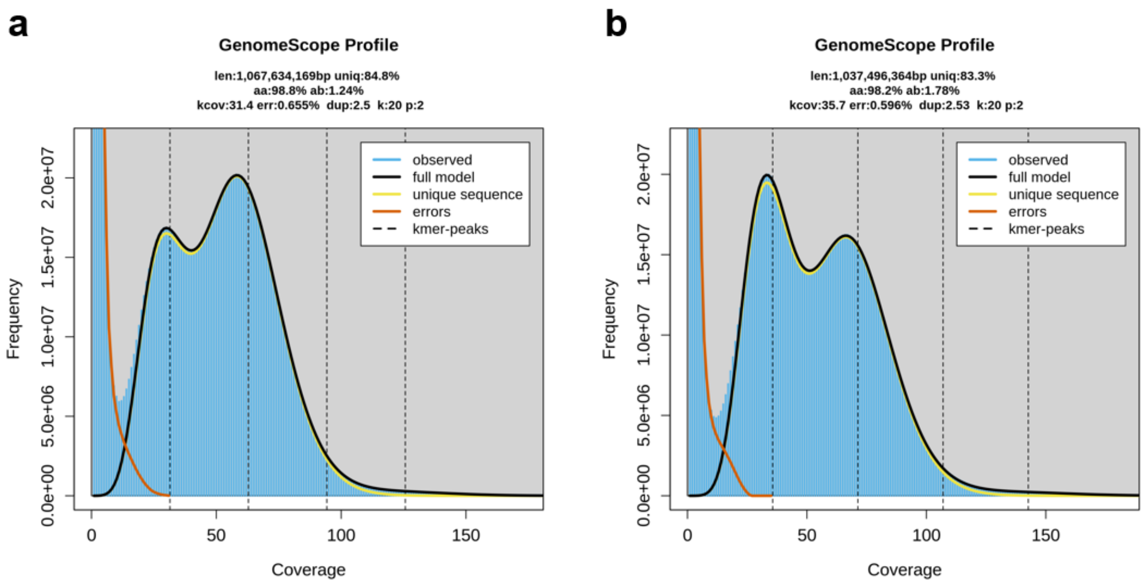


**Figure S4.** Genomescope profile of zebra finch assemblies calculated from 10X-Linked reads of. **a.** bTaeGut1.4 assembly and **b.** bTaeGut2 assembly.

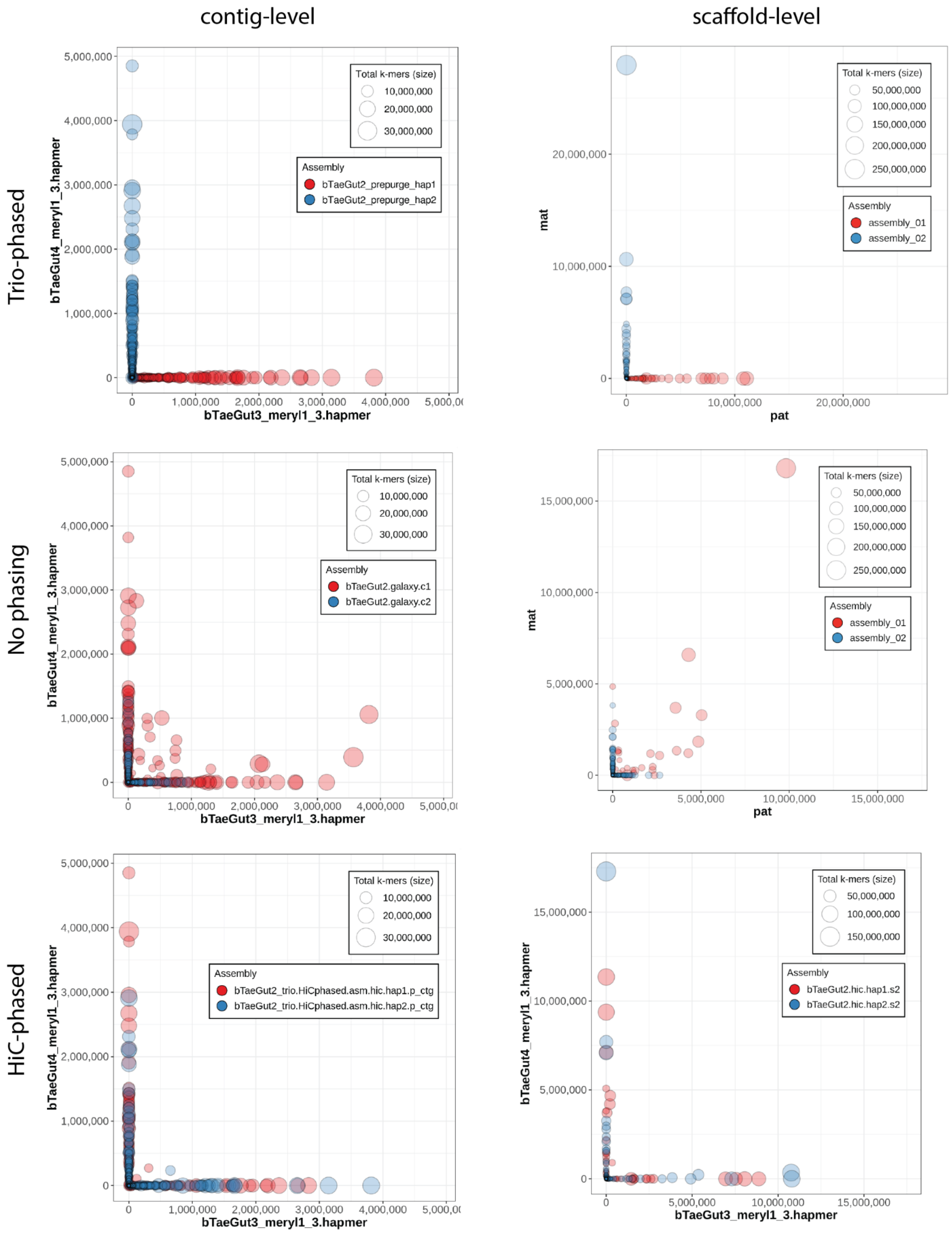


**Figure S5**. Merqury blob plots for *Taeniopygia guttata* (bTaeGut2) contigs with various hifiasm modes: top, trio-phased assembly; middle, pseudohaploid ("solo") assembly; and bottom, HiC-phased assembly. Each circle represents a contig, and the size of the circle corresponds to the overall size of the contig, while the color of the circle represents which assembly the contig is in, and the position of the circle along the X- and Y-axes corresponds to parental hapmer content. Contigs that are considered to be properly phased (e.g., no switch errors) will be flush with either the X- or Y-axis depending on which parental haplotype the contig corresponds to, since this means that the contig contains a number of hapmers from one parent and none from the other parent. This is illustrated in the trio-phased plots (top row), where each contig has been properly phased using parental data. The pseudohaplotype contigs/scaffolds (middle row) shows numerous sequences with a mix of parental hapmer content, as they are not flush to either of the axes. The Hi-C-phased (bottom row) sequences are largely properly phased, with a few contigs off the axes.

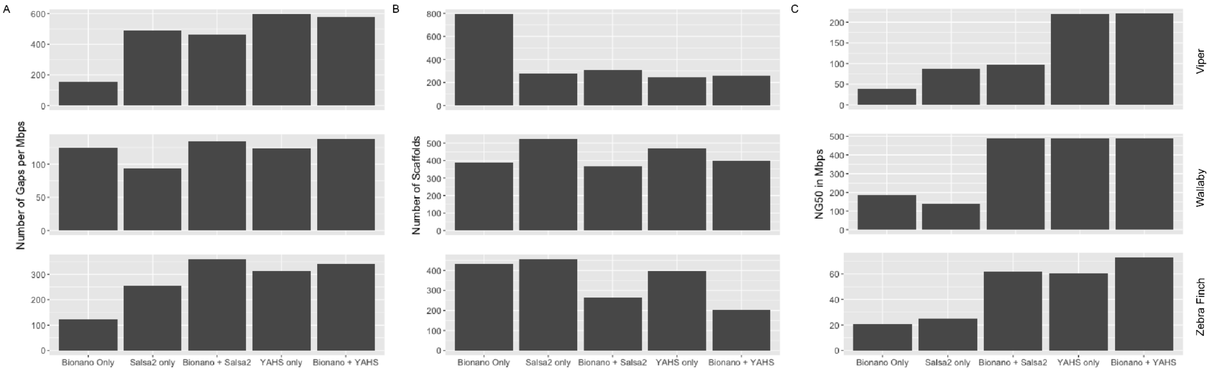


**Figure S6.** Comparison of scaffolding methods. Comparison of assembly quality between Bionano scaffolding only, Salsa scaffolding only, Bionano and Salsa Scaffolding, Yahs Scaffolding only, and Bionano and Yahs scaffolding. The comparison has been made for the Viper, the Wallaby, and the Zebra Finch Assemblies. **a.** Number of gaps per Mbps; **b.** Number of Scaffolds; **c**. NG50 in Mbp

**A**

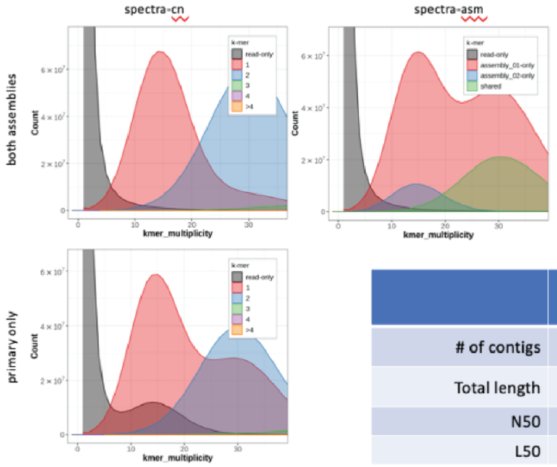


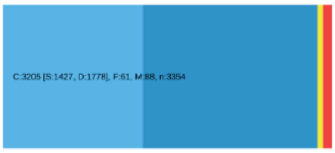


| | Primary (unpurged) | Alternate (unpurged) |
|---|---|---|
| # of contigs | 3,548 | 3,805 |
| Total length | 7,012,181,570 | 1,303,392,418 |
| N50 | 5,385,022 | 1,773,313 |
| L50 | 358 | 208 |

**B**

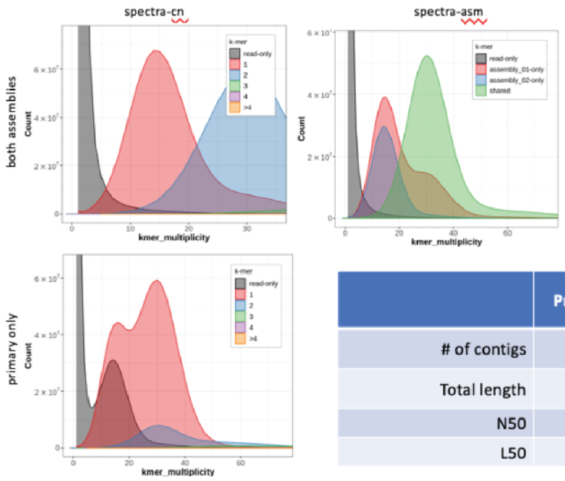


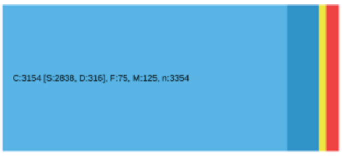


| | Primary (post-purging) | Alternate (post-purging) |
|---|---|---|
| # of contigs | 1,363 | 3,997 |
| Total length | 4,663,182,711 | 3,473,507,201 |
| N50 | 7,405,859 | 2,218,743 |
| L50 | 184 | 444 |

**C**

| | Hap1 | Hap2 |
|---|---|---|
| # of contigs | 2,511 | 2,172 |
| Total length | 4,339,321,113 | 4,302,114,182 |
| N50 | 5,289,116 | 5,035,338 |
| L50 | 232 | 240 |

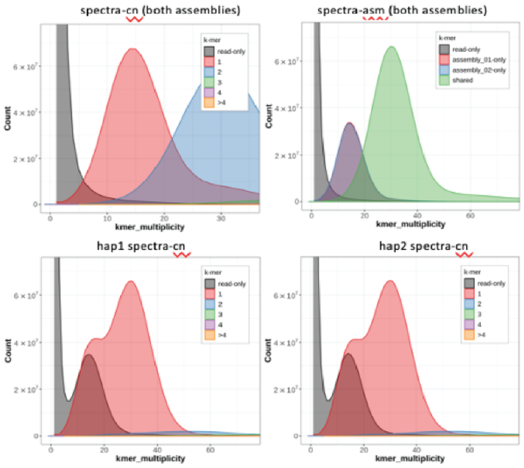


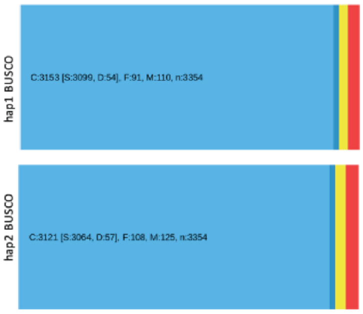


**Figure S7.** Quality control for several *Gastrophryne carolinensis* (aGasCar1) assemblies: **a.** pseudohaplotype assembly pre-purging, **b.** pseudohaplotype post-purging, and **c.** Hi-C-phased contigs. The quality control metrics shown are merqury *k*-mer spectra graphs, assembly statistics, and BUSCO genes for Vertebrata. The merqury plots show the partitioning of *k*-mers from the readset across the two assemblies, which can signal that the two assemblies are unbalanced such as in S8b, where there are regions with diploid coverage present only in the primary assembly (spectra-asm plot in panel B), and at 2-copy (seen in the primary-only merqury plot in panel B and the BUSCO image for the same panel). Panel C shows the spectra-cn plot for hap1 and hap2 individually, and there are much fewer 2-copy *k*-mers at diploid coverage.

The pre-purging pseudohaplotype assemblies show that the assemblies are largely unbalanced, with many diploid regions being retained twice in the primary assembly. Purging addresses this, but unsatisfactorily, as there are still a significant amount of BUSCO duplicates present as well as unevenness between the two haplotypes. In contrast, the Hi-C-phased contigs show proper haplotype resolution from the start, with minimal BUSCO duplicates and the two haplotypes resolved on a *k*-mer level from the start.
